# Mutations within and outside the PfFNT transport pathway drive MMV007839 resistance in *Plasmodium falciparum*

**DOI:** 10.64898/2026.07.08.737153

**Authors:** Ciara Wallis, Abigail Partridge, Adele Lehane, Ben Corry

**Author notes:** These authors contributed equally (Ciara Wallis, Abigail Partridge).

## Abstract

*Plasmodium falciparum*, the parasite responsible for the majority of malaria cases and deaths worldwide, poses a major global health challenge due to its ability to rapidly develop resistance to antimalarial drugs. To circumvent existing resistance pathways, it is of interest to identify both new inhibitors and new therapeutic targets within the parasite. One such target is the *Plasmodium falciparum* Formate–Nitrite Transporter (PfFNT), which mediates essential lactate and proton (H^+^) export from the parasite cytosol during the intra-erythrocytic stage. Several inhibitors of PfFNT have been identified that block lactate/H transport and/or kill parasites *in vitro*, including MMV007839, which binds within the substrate transport pathway as revealed by cryo-EM structures. However, three resistance-conferring mutations have previously been identified following prolonged *in vitro* exposure of parasites to PfFNT inhibitors. Here, we performed further *in vitro* evolution experiments with MMV007839 in *P. falciparum* and identified two additional mutations in PfFNT, V200L and L198M, the latter of which is located outside the substrate transport pathway. These mutations are associated with 6-fold and 13-fold changes in the IC_50_ of MMV007839. Using molecular dynamics simulations, and thermodynamic integration, we show that these mutations, together with previously identified resistance mutations, destabilise inhibitor binding within the transport pathway; and we suggest the molecular mechanisms underlying this destabilisation. Finally, using both molecular docking and additional simulations, we suggest new sites where additional mutations that confer resistance to PfFNT inhibitors may arise.

**Author Summary:** Malaria parasites are becoming resistant to existing drugs, creating an urgent need for new treatments. One promising drug target is the *Plasmodium falciparum* Formate-Nitrite Transporter (PfFNT), which the parasite needs to remove waste products and survive during infection. Although some compounds that block PfFNT have been identified, several mutations in the protein can make these compounds ineffective. To better understand how resistance might develop, we exposed malaria parasites to the PfFNT inhibitor MMV007839 in the laboratory and identified two new resistance-related mutations, one of which lies outside the MMV007839-binding site. We also used computer models to predict other regions where resistance-conferring mutations could occur. These findings help assess PfFNT as a drug target and provide guidance for designing new drugs that are less likely to be undermined by resistance.

## Introduction

Malaria is a mosquito-transmitted disease that, in 2024, was associated with an estimated 282 million cases and 610,000 deaths.^1^ The disease is caused by six species of *Plasmodium* parasites; however, most cases and deaths are caused by the species *Plasmodium falciparum.*^1,2^ Control of malaria is heavily dependent on both preventing mosquito transmission and being able to effectively treat infections with antimalarials.^1,3^ However, the ability of *P. falciparum* to develop resistance to almost every clinically available antimalarial is challenging treatment efforts.^4^ Emergence of resistance to artemisinin-based combination therapies, the current frontline therapies, in Southeast Asia, South America, and Africa highlights a need for the identification of both new antimalarial drugs and new antimalarial drug targets.^5–9^ Parasite targets that have not previously been exploited by clinically available antimalarials are particularly attractive, as compounds that act on these pathways will be less likely to be susceptible to existing resistance mechanisms.^10^

One potential target is the *P. falciparum* Formate–Nitrite Transporter (PfFNT), an essential protein localised to the parasite plasma membrane during the disease-causing stage of its lifecycle.^11,12^ PfFNT mediates the export of lactate and H^+^, the major by-products of parasite energy metabolism during this stage (**Fig. 1a**), preventing cytosolic acidification, cell swelling, and parasite death occurring from the build-up of these metabolites within the cytosol.^11,12^ In 2017, the compounds MMV007839 and MMV000972, identified from a screen of the Medicines for Malaria Venture (MMV) “Malaria Box”, were shown to potently inhibit PfFNT-mediated [^14^C]-lactate transport in *P. falciparum* parasites as well as in *Saccharomyces cerevisiae* and *Xenopus laevis* oocytes expressing PfFNT.^13,14^ These compounds also inhibited parasite proliferation *in vitro*, with 50% growth inhibitory concentrations (IC_50_) values of 140 nM for MMV007839 and ∼1.8 μM for MMV000972.^13,14^ Furthermore, the compounds have been shown to induce acidification of the parasite cytosol, swelling of both isolated parasites and parasitised erythrocytes, and accumulation of lactate within parasitised erythrocytes.^13^ However, due to findings of toxicity in some assays, including in assays with human embryonic kidney (HEK293) cells^15^, human liver (HepG2) cells^15,16^, and zebrafish embryos^16^, these compounds are unsuitable for clinical development. Many other PfFNT inhibitors have since been synthesised, with many sharing a common pentafluoro-3-hydroxy-pent-2-en-1-one scaffold. Among these are compounds that display micromolar (BH296, BH297) or submicromolar (BH267.meta) antiplasmodial activity and, importantly, negligible toxicity (BH267.meta) in mouse models.^14,15,17,18^

**Figure 1.**
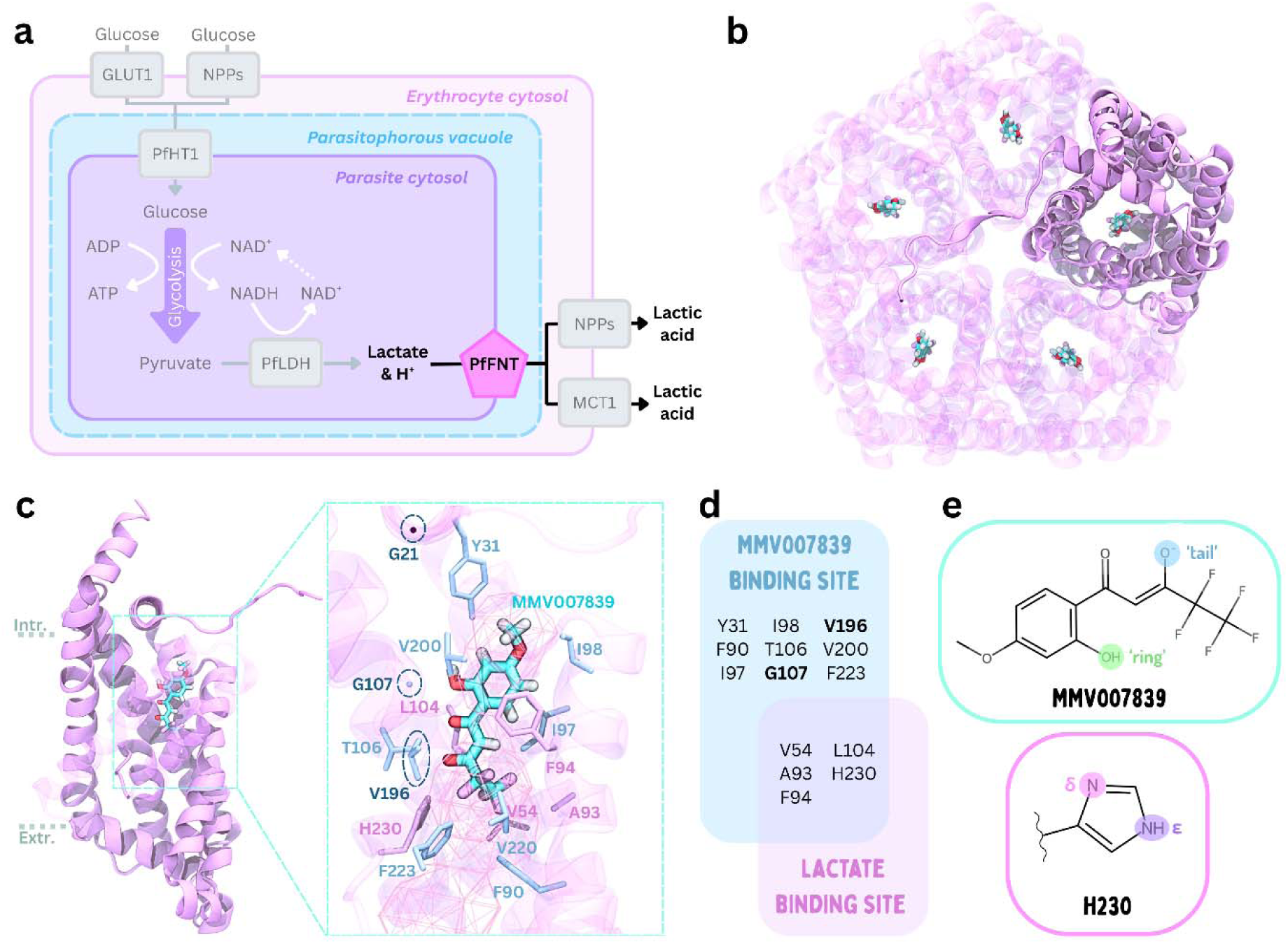
Comparisons between the MMV007839 and lactate binding sites suggests potential sites for resistance conferring mutations. **a)** Schematic of PfFNT’s role as a lactate/H^+^ exporter in parasite energy metabolism during the intraerythrocytic stage. Other proteins, such as the human (GLUT1)^27^ and parasite (PfHT1)^28,29^ glucose transporters, the parasite lactate dehydrogenase enzyme (PfLDH)^30^, and the human lactate exporter (MCT1)^31^ are shown together with the parasite-induced new permeabilit pathways^32,33^ that help facilitate nutrient uptake and export. **b)** Cryo-EM structures of PfFNT resolved with MMV007839 show the inhibitor bound in each subunit (PDB ID: 7E27). PfFNT is shown in a pin cartoon representation, with one of the five subunits highlighted. **c)** A close up of the transport cavit viewed from the plane of the membrane shows residues that contact both lactate^26^ and MMV007839^19,20^ (pink) or MMV007839 only (blue). Residues implicated as sites for resistance mutations are highlighted by dashed circles. The mesh represents a volumetric surface of the pathway taken by lactate/lactic acid through PfFNT. **d)** Venn diagram highlighting the overlap between the residues forming the MMV007839 and lactate binding sites. Sites of known resistance conferring mutations are bolded. **e)** Structures of linear MMV007839 (top) and H230 (bottom). MMV007839 contains two hydroxyl groups: a phenolic “ring” OH (green) and an aliphatic “tail” OH (blue). H230 contains two protonatable nitrogens in its side chain: the δ-nitrogen (pink) and the ε-nitrogen (purple). Only the side chain of H230 is shown; the remainder of the residue is omitted and indicated by the wavy line.

Prolonged exposure of *P. falciparum* parasites to either MMV007839 or BH267.meta *in vitro* resulted in the identification of three mutations associated with resistance: G21E, G107S, and V196L.^13,14,17^ In 2021, five PfFNT structures were solved in either the apo state or with MMV007839 bound within the cavity.^19,20^ The structures reveal the protein to have a homopentameric structure characteristic of the FNT family^21–25^ (**Fig. 1b**), with each subunit containing a transport cavity bordered by hydrophobic constrictions formed by F94, I97, and L104 on the intracellular side and F90, F223, and H230 on the extracellular side. The structures with MMV007839 bound reveal that the inhibitor binds in its linear tautomeric form within each transport cavity (**Fig. 1b-c,e**).^19,20^ Although structures of PfFNT have not been resolved with other known inhibitors bound, all potent PfFNT inhibitors share a common chemical scaffold, suggesting these analogues are likely to bind in a similar location within the cavity as MMV007839. Therefore, the MMV007839-bound PfFNT structures can be used as a useful model for predicting how mutations affect the binding of MMV007839 and other chemically similar compounds.

Although cryo-EM structures of PfFNT were not resolved with lactate bound, previous molecular dynamics studies of PfFNT indicate that the lactate binding site overlaps substantially with the MMV007839 binding site.^26^ Notably, several residues that contact MMV007839 do not appear to be essential for lactate binding, including V196 and G107 - sites of previously identified resistance mutations (**Fig. 1c-d**).^13,14,17,26^ While some studies have suggested that the narrow PfFNT cavity limits mutational flexibility^14,17^, the appearance of the G21E mutation in MMV007839-selected parasites^17^, located within the substrate transport pathway but outside the lactate binding site, suggests other mutations may arise in sites that are not essential for substrate transport.

In this study, we examined how the previously identified G107S, V196L, and G21E mutations destabilise MMV007839 binding, and investigated the possibility of other resistance conferring mutations arising in PfFNT using *in vitro* evolution experiments with *P. falciparum* parasites, together with molecular dynamics simulations and molecular docking. We identified a further two novel mutations in PfFNT, V200L and L198M, the latter of which is located outside of the inhibitor binding site. Both mutations desensitise parasites to MMV007839 without causing an obvious growth defect. Using thermodynamic integration simulations, we show that these mutations destabilise MMV007839 binding by altering the flexibility of helices bordering the transport pathway. In ‘flooding’ simulations, in which we include a high concentration of lactate molecules to improve sampling of possible binding sites, we show lactate is still able to bind within the same site we previously identified for wildtype PfFNT.^26^ This supports our experimental observations that the PfFNT-V200L and PfFNT-L198M parasites maintain a robust growth rate. As our data indicate that several mutations in the PfFNT transport pathway can confer resistance to MMV007839, we performed a large-scale molecular docking screen alongside thermodynamic integration simulations to identify additional sites that may accommodate resistance-conferring mutations. Of the mutations chosen for further study with thermodynamic integration, we find that mutation of Valine to Leucine for residues 54, 203, 216, and 220 destabilises MMV007839 binding without preventing the binding of lactate in flooding simulations. As many PfFNT inhibitors share similar scaffolds, it is possible that some of the mutations that we and others have identified or predicted in PfFNT could confer cross-resistance. This information can be applied to aid in assessing the worth of PfFNT as a drug target and improve the rational design of PfFNT inhibitors to avoid likely resistance-conferring mutations.

## Results

### Mutations associated with resistance to MMV007839 destabilise its binding within the transport cavity

As MMV007839 is the only PfFNT inhibitor for which a binding site has been resolved experimentally, we first sought to define whether the resistance-conferring mutations G21E, G107S, and V196L alter MMV007839’s binding within the transport pathway. While previous studies have used molecular docking to investigate the effects of the G107S and V196L mutations on MMV007839 binding^17,19^, the effect of the G21E mutation had only been hypothesized. Further, as docking only involves a static structure, it was not clear whether there may be additional conformational changes that occur following introduction of these mutations.

To investigate this, we performed triplicate 1 μs molecular dynamics simulations starting from the PfFNT-WT structure with MMV007839 bound within the cavity, and computationally introduced each of the known mutations associated with resistance to MMV007839 (**Fig. 2a**). In each mutant system, MMV007839 remained stably bound within the cavity over the course of the simulation as indicated by minimal changes in the position of the inhibitor within the cavity across the simulation (**Fig. S1a-b**). To determine whether MMV007839 binding was destabilised in any of the mutant systems, we performed thermodynamic integration simulations to calculate the relative change in binding free energy (ΔΔG) of MMV007839 in the mutant systems relative to the wildtype systems. In all mutant systems, we found that MMV007839 binding was destabilised, as shown by the positive ΔΔG values of 18.0 ± 0.7 kJ/mol, 18.3 ± 1.1 kJ/mol, and 10.4 ± 1.1 kJ/mol for PfFNT-G107S, PfFNT-V196L, and PfFNT-G21E, respectively (**Fig. 2b**, **Table S4**).

**Figure 2.**
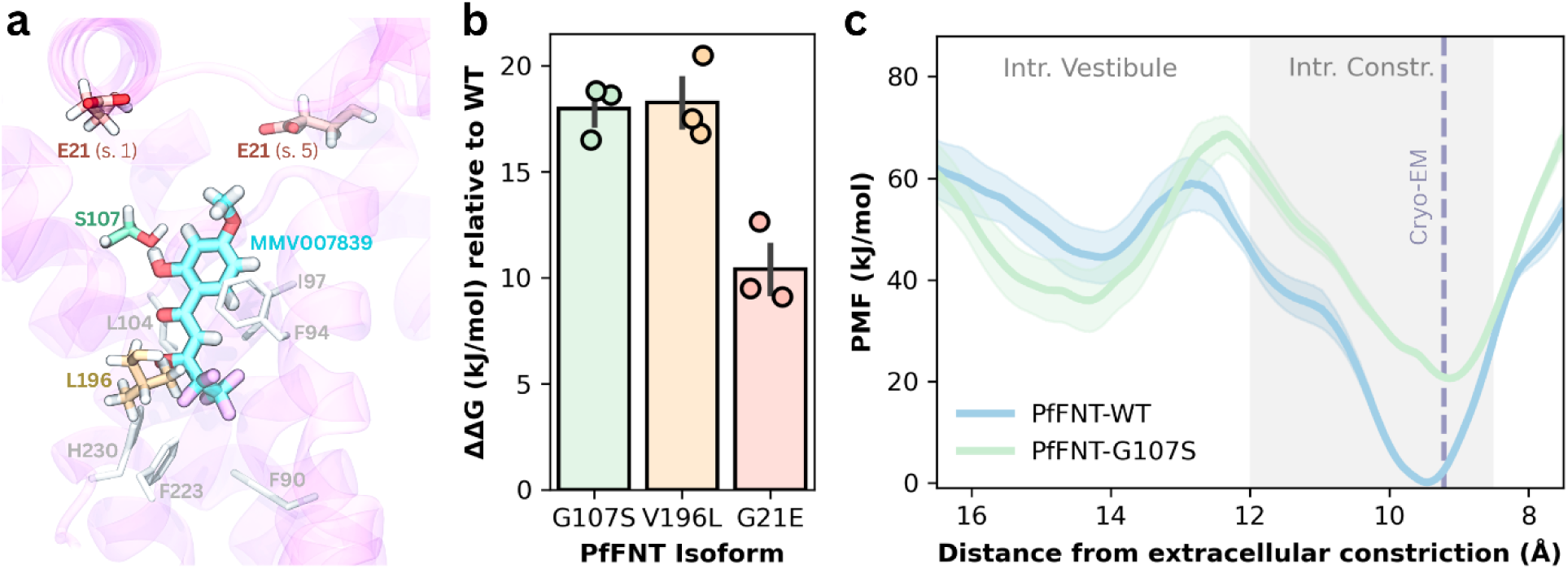
The G21E, G107S, and V196L mutations destabilise MMV007839 binding. **a)** Computationally generated mutant structures of PfFNT-G21E, PfFNT-G107S, and PfFNT-V196L were aligned to show the location of each mutation relative to the MMV007839 binding site in the PfFNT-WT cryo-EM structure. The sidechains of S107 (green), and L196 (orange) are shown in licorice representation. Sidechains of E21 from subunit 1 (s. 1) and subunit 5 (s. 5) are shown in red. Intracellular (F94, I97, L104) and extracellular (F90, F223, H230) constriction residues are also shown in a grey licorice for reference. **b)** Mean change in binding free energy (ΔΔG) for MMV007839 binding in the cavity in the PfFNT-G107S (green), PfFNT-V196L (orange) and PfFNT-G21E (red) systems, relative to binding in the cavity in the PfFNT-WT systems. Averages were obtained from three independent sets of TI simulations. Error bars = SEM. **c)** Average potential of mean force (PMF) (kJ/mol), ± SEM, *n* = 5, for the movement of MMV007839 into the cavity in PfFNT-WT (blue) and PfFNT-G107S (green) systems from one-dimensional umbrella sampling simulations. The distance between the centre of mass (COM) of MMV007839 and the COM of the extracellular constriction is represented along the x-axis. The shaded grey region indicates the location of the intracellular constriction (‘Intr. Constr.’). The dashed purple line i used to represent the distance between the COM of MMV007839 and the COM of the extracellular constriction in the 7E27 cryo-EM structure (‘Cryo-EM’). The location of the intracellular vestibule above the constriction is also shown (‘Intr. Vestibule’).

As the thermodynamic integration simulations indicate binding is heavily destabilised but the inhibitor does not leave the cavity in our standard MD simulations, we next investigated whether the G107S mutation alters the ability of the inhibitor to enter or leave the binding site using umbrella sampling simulations. In PfFNT-WT, MMV007839 has a small barrier to pass the intracellular constriction formed by F94, I97, and L104 and has a deep energy minimum within the cavity. Comparatively, MMV007839 still has a deep minimum within the cavity in the PfFNT-G107S system but has a larger barrier for passing the constriction to enter. A second, smaller minimum can also be observed higher up along the transport pathway, which appears to correspond to the hydroxyl group near the fluoroalkyl tail of MMV007839 interacting with the hydroxyl group on S107 (**Fig. 2c**).

To further characterise how inhibitor–residue interactions are perturbed in each mutant relative to the wild-type system, per-residue Molecular Mechanics/Generalized Born Surface Area (MM/GBSA) residue decomposition analysis was performed. In PfFNT-G107S, interactions between the inhibitor and residues 103–108, as well as Y31 and V196, become destabilised. In PfFNT-V196L, the strongest effect is observed for T106, while in PfFNT-G21E, interactions between MMV007839 and residues I97 and I98 become destabilised (**Fig. S1c**).

Additionally, introduction of the negatively charged glutamate in PfFNT-G21E appears to alter the protein’s salt-bridge network. In the wild-type protein, a stable salt bridge is maintained between K34 and E22 throughout most of the simulation; however, in the G21E mutant, E21 replaces E22 as the primary interaction partner of K34 (**Fig. S1d**). We propose that this disruption of the salt-bridge network subtly reshapes the binding pocket, contributing to destabilisation of MMV007839 binding.

As the G21E mutation occurs outside of the cavity and there are other residues in the MMV007839 binding site that do not directly overlap with the lactate binding site, we investigated the possibility of other resistance mutations being able to occur in PfFNT. Using *in vitro* evolution experiments, we exposed 10 *P. falciparum* cultures (Dd2^PfFNT-WT^), each containing ∼4 x 10^8^ parasites, to 750 nM MMV007839 (approximately 3-fold its IC_50_ value for growth inhibition) (**Fig. S2a**). After 12-14 days, parasite proliferation was detected in six flasks (**Fig. S2b**). DNA was extracted from parasites from the six flasks and the *pffnt* gene was sequenced. This resulted in the identification of two novel mutations; a T>A base change at position 592 of *pffnt*, resulting in a Leu198 to Met198 amino acid change (L198M) was found in parasites from one flask, and a G>T base change at position 598 of *pffnt*, resulting in a Val200 to Leu200 amino acid change (V200L) was found in parasites from five flasks (**Fig. S2c**). No resistant parasites were observed in the remaining four flasks within the 52 days of the experiment.

We next measured the sensitivity of the mutant parasites to growth inhibition by MMV007839. The Dd2^PfFNT-L198M^ and Dd2^PfFNT-V200L^ parasites were significantly less sensitive to MMV007839 than their Dd2^PfFNT-WT^ parents with 6.6 ± 0.2-fold and 13.7 ± 0.5-fold higher IC_50_ values, respectively, but were not as resistant to the inhibitor as Dd2^PfFNT-G107S^ mutant parasites (generated previously using the same parental line^13^), for which a 235 ± 26-fold elevated IC_50_ was observed (generated previously using the same parental line^13^) (**Fig. 3a**, **Fig. S2e, Table S1**). The IC_50_ values for MMV007839 obtained here for Dd2^PfFNT-WT^ and Dd2^PfFNT-G107S^ parasites (0.28 ± 0.02 µM and 65 ± 5 µM) were within ∼2-fold of those reported previously for the same parasites^13^. As expected, none of the PfFNT mutations affected parasite sensitivity to the antimalarial chloroquine (**Fig. S2d,f, Table S2**). Parasite proliferation assays were also performed for the five flasks of parasite cultures in which the V200L mutation was found. The Dd2^PfFNT-V200L^ parasites originating from the five different flasks did not differ from each other in their responses to MMV007839 or chloroquine (**Fig. S2g-h**). Flask #4 was used as a representative V200L culture for all subsequent experiments.

**Figure 3.**
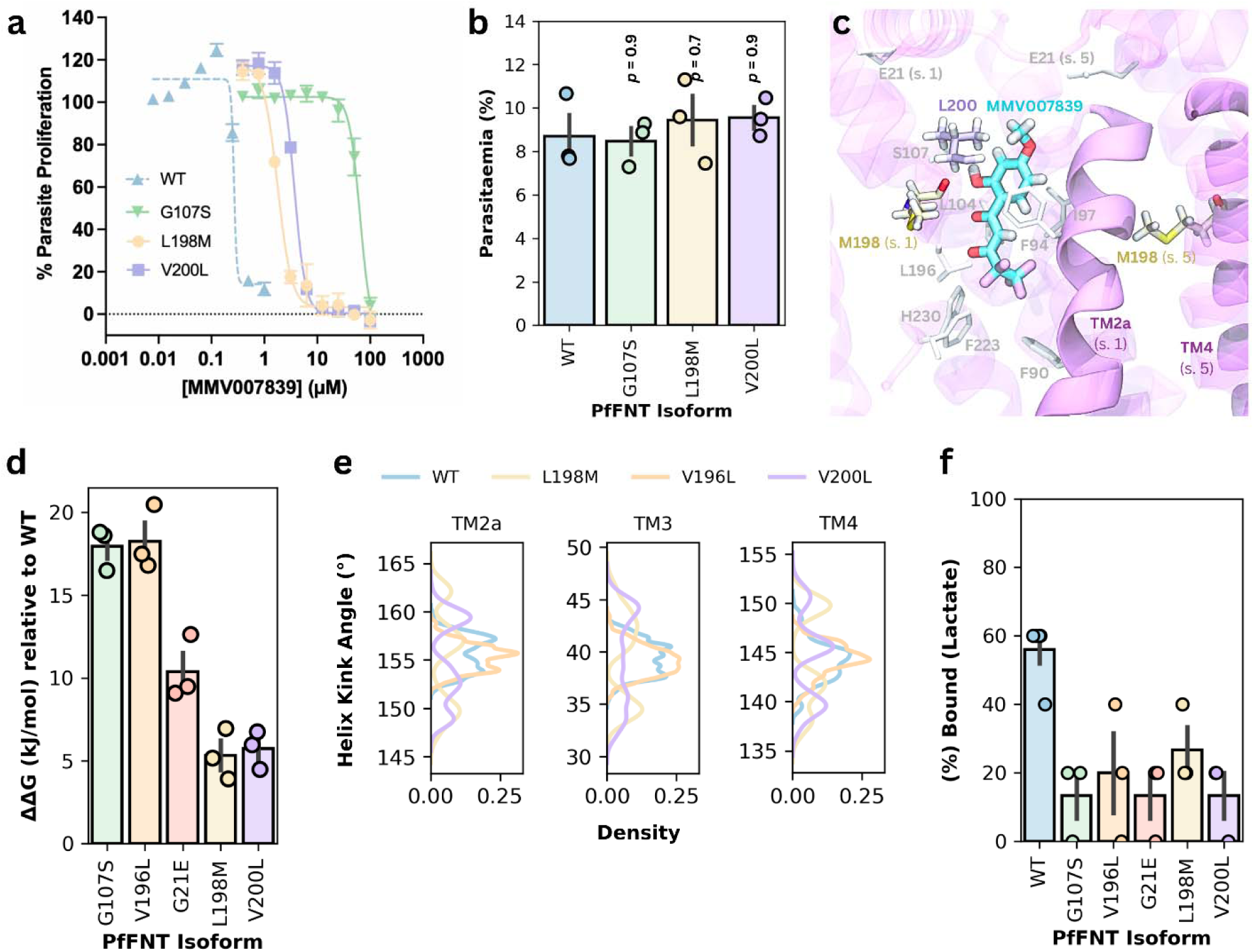
Mutations associated with resistance to MMV007839 can occur within or outside the transport pathway. The L198M and V200L mutations discovered in this study from *in vitro* evolution experiments **a)** desensitise parasites to MMV007839 exposure in parasite proliferation assays, **b)** without appearing to give rise to a major growth defect. The mean sensitivities of wildtype (blue, upward triangles), G107S (green, downwards triangles), L198M (yellow, circles), and V200L (purple, squares) cultures to MMV007839 are plotted (**a**). Parasite growth was assessed by calculating parasitaemia 48 h after adjusting parasite cultures to a parasitaemia of 0.8-1% (**b**). Bars represent the mean parasitaemia for wildtype (blue), G107S (green), L198M (yellow), and V200L (purple) parasite cultures. Symbols show the data from each independent experiment. Data for parasite proliferation assays and parasite growth comparisons were obtained from three experiments (with the exception of the highest concentration for Dd2^PfFNT-WT^ in (**a**), which was n = 2; containing internal triplicates in the case of the proliferation assays) each performed on separate days. Error bars for **a** and **b** are SEM. Statistical significance was determined using one-way ANOVA with a *post-hoc* Tukey test. IC_50_ values are reported in **Figure S2** and **Table S1**. **c)** Computationally generated mutant structures of PfFNT-L198M and PfFNT-V200L were aligned to show the location of each mutation relative to the MMV007839 binding site in the PfFNT-WT cryo-EM structure. The sidechains of V200L (purple) in subunit 1 and L198M (yellow) in subunits 1 and 5 (s. 5) are shown. The sidechains of E21, S107, L196, and the intracellular (F94, I97, L104) and extracellular (F90, F223, H230) constriction residues, are also shown in a grey licorice for reference. TM2a on subunit 1 (s. 1) and TM4 on subunit 5 (s. 5) are shown in solid pink cartoon, while the rest of the protein is shown in transparent cartoon. **d)** Calculations of the average change in binding free energ (ΔΔG) for MMV007839 binding in the cavity in the PfFNT-L198M (yellow) and PfFNT-V200L (purple) systems, relative to PfFNT-WT, from TI simulations. The ΔΔG values for the PfFNT-G107S (green), PfFNT-V196L (orange), PfFNT-G21E (red) systems are the same as those shown in **Fig. 2b** and are included here for comparison. Data are from three independent sets of TI simulations. Error bars = SEM. **e)** Density plots of the kink angles (°) adopted by the TM2a, TM3, and TM4 helices from PfFNT-WT (blue), PfFNT-V196L (orange), PfFNT-L198M (yellow), and PfFNT-V200L (purple) simulations. Data were combined from each subunit from three replicates across 1 µs of simulation per replicate. **f)** Average percentage of subunits bound by lactate from flooding simulations out of the total number of binding sites available for simulations of the PfFNT-WT (blue), PfFNT-G107S (green), PfFNT-V196L (orange), PfFNT-G21E (red), PfFNT-L198M (yellow), and PfFNT-V200L (purple) systems. Mutant simulations were performed for 1 µs per replicate for three independent replicates. Error bars = SEM. PfFNT-WT simulations had been run previously for 2 µs per replicate for eight independent replicates. Statistical significance was assessed using a binomial generalized linear model (GLM) with a logit link function, with mutation treated as a fixed effect, followed by Dunnett-adjusted post hoc comparisons against WT. The adjusted *p* values for all comparisons are reported in **Table S6**.

PfFNT is essential for parasite survival^13,14,17^; however, it has previously been shown from [^14^C]-lactate uptake experiments in parasites, PfFNT-expressing *X. laevis* oocytes, and PfFNT-expressing *S. cerevisiae* that the G107S mutation in PfFNT causes ∼30-60% reduction in lactate transport.^13,14^ Despite this reported reduction in transport, no detectable growth defect was observed during the asexual *P. falciparum* blood stage for G107S cultures.^13^ To investigate whether the L198M or V200L mutations in PfFNT might impose a fitness cost, we performed a preliminary short-term comparison of the growth of Dd2^PfFNT-L198M^, Dd2^PfFNT-V200L^, Dd2^PfFNT-WT^ and Dd2^PfFNT-G107S^ parasites. Cultures were prepared with a haematocrit of 2% and a parasitaemia of 0.8-1%, and the parasitemia was determined again after 48 hours. From these experiments, no significant differences in growth were observed between the PfFNT-mutant and parental parasites (**Fig 3b**).

To better understand how the L198M and V200L mutations may confer resistance to MMV007839, we examined where they are found on the protein (**Fig. 3c**). L198M and V200L both occur on TM4, the same TM helix as the V196L mutation. Like V196L, V200L is located inside the transport cavity, while L198M is located at the interface between subunits. Estimates of ΔΔG from TI simulations suggest that the L198M and V200L mutations destabilise MMV007839 binding, but to a lesser degree than the G107S, V196L, and G21E mutations (**Fig. 3d**). MM/GBSA calculations for the L198M system indicate that interactions between MMV007839 and residues on the N-terminal helix (NTH) and transmembrane (TM) helices TM2a, TM2b, TM3, TM4, and TM5a get destabilized following introduction of the mutation (**Fig. S1d**). Further analysis of the kink angles adopted by all helices within the PfFNT subunit across the simulation reveals that in both the V200L and L198M systems, TM1, TM2a, TM2b, TM3, TM4, and TM5a all become more flexible relative to the wildtype and V196L systems (**Fig 3e, Fig. S3a**). In addition to the increased flexibility of these TM helices, measurements of the transport pathway radius from the V200L and L198M systems show slightly narrower profiles than in the wildtype PfFNT system (**Fig. S3b**).

To predict whether lactate binding and transport were likely to be impaired by any of these mutations, we next performed computational flooding simulations with lactate, similar to what was previously performed to investigate the transport mechanism of PfFNT-WT.^26^ At least one lactate binding event could be observed in each simulation system, although the number of lactate binding events appeared reduced in comparison to the wildtype system (**Fig. 3f**). This suggests that, although these mutations may slightly alter the shape and dynamics of the transport cavity, PfFNT is likely to remain functional as a lactate transporter.

Having identified two additional mutations associated with MMV007839 resistance in PfFNT, we wanted to investigate the possibility of other mutations occurring along the transport pathway that could destabilise MMV007839 binding. As it may be challenging to identify all possible resistance-conferring mutations using in vitro evolution experiments, we performed a molecular docking screen of MMV007839 to structures harbouring all possible single amino acid substitutions along the transport pathway, including mutations to lactate binding residues, and compared the position of MMV007839 in the top clustered binding pose in each mutant to its position in the WT cryo-EM structure. Mutations were classified as likely to be destabilising if the root mean square deviation (RMSD) of the docked MMV007839 pose was greater than 3.5 Å from the inhibitor’s position in the 7E27 structure. The cutoff of 3.5 Å was determined by plotting the distribution of RMSD values from all mutant docked poses (**Fig. S4c**). From docking results for the 1560 mutations, we filtered for mutations that required only a single base change and had a docked pose RMSD greater than 3.5 Å (**Fig. 4a**). Of the five known mutations associated with resistance to MMV007839, only G107S was identified out of the remaining 110 mutations. Docking to other mutant structures yielded similar poses to docking to PfFNT-WT (**Fig. S4a-e**).

**Figure 4.**
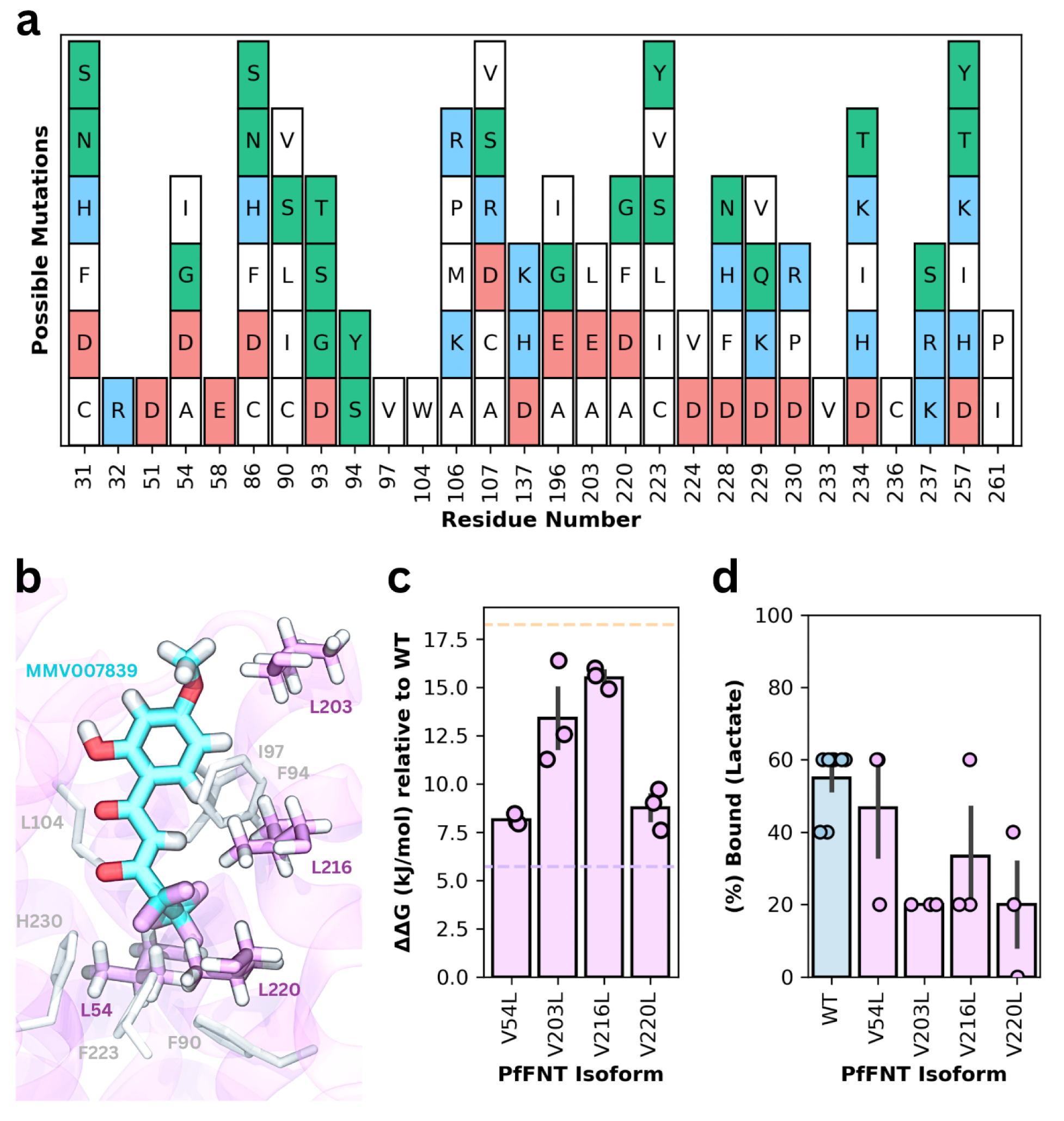
Additional mutations may be possible that destabilise MMV007839 binding without inhibiting lactate passage. **a)** Potential mutations identified from a docking screen after filtering for mutations that only require a single base change and in which the top cluster docked pose had an RMSD greater than 3.5 Å from the position of MMV007839 in its cryo-EM binding site. Bars are coloured by mutant sidechain. Non-polar residues are shown in white, basic residues are shown in blue, acidic residues are shown in red, and polar residues are shown in green. **b)** Computationally generated mutant structures of PfFNT-V54L, PfFNT-V203L, PfFNT-V216L, and PfFNT-V220L were aligned to show the location of each mutation relative to the MMV007839 binding site in the PfFNT-WT cryo-EM structure. Mutant sidechains are shown in pink licorice. The sidechains of the intracellular (F94, I97, L104) and extracellular (F90, F223, H230) constriction residues, are also shown in a grey licorice for reference. **c)** Calculations of the average change in binding free energy (ΔΔG) for MMV007839 binding in the cavity in PfFNT-V54L, PfFNT-V203L, PfFNT-V216L, and PfFNT-V220L systems, relative to PfFNT-WT, from TI simulations. Data are from three independent sets of TI simulations. Error bars = SEM. Dashed lines showing ΔΔG for PfFNT-V196L (orange) and PfFNT-V200L (purple) are added as a reference. **d)** Average percentage of subunits bound by lactate from flooding simulations out of the total number of binding sites available for simulations of the PfFNT-WT (blue) and PfFNT-V54L, PfFNT-V203L, PfFNT-V216L, and PfFNT-V220L (pink) systems. Mutant simulations were performed for 1 µs per replicate for three independent replicates. Bars for **c** and **d** represent mean values, individual datapoints are shown as points on each bar, and error bars represent SEM. PfFNT-WT simulations had been run previously for 2 µs per replicate for eight independent replicates^26^. Statistical significance was assessed using a binomial generalized linear model (GLM) with a logit link function, with mutation treated as a fixed effect, followed by Dunnett-adjusted post hoc comparisons against WT. The adjusted *p* values for all comparisons are reported in **Table S6**.

We also applied our docking pipeline to two other recently identified PfFNT inhibitors – Compounds 7e and 8a.^34^ These inhibitors were shown to have IC_50_ values for wildtype 3D7 parasites of 14.8 ± 0.9 nM and 49.8 ± 0.5 nM, respectively.^34^ Multiple mutations were suggested by docking to destabilise the binding of all three inhibitors, including F90I, F90L, I97V, V196I, F223I, F223L, and N234T (**Fig. S5a-c**). If any of these mutations should arise from prolonged exposure of parasites to these inhibitors, it is important to consider the possibility of cross-resistance occurring due to the inhibitors sharing a common scaffold.

Two of the five mutations previously associated with resistance to MMV007839 involve a Valine to Leucine substitution. In addition to V196 and V200, four other valine residues line the PfFNT transport pathway – V54, V203, V216, and V220. In the related FNTs from *Toxoplasma gondii* (TgFNT1-3), the residue equivalent to V54 is an isoleucine in TgFNT1, the residue equivalent to V216 is a leucine in TgFNT1 and a methionine in TgFNT2 and TgFNT3, and the residue equivalent to V220 is an isoleucine in all three TgFNTs. In contrast, the positions corresponding to V196, V200, and V203 are conserved as valines in TgFNT1-3 (**Fig. S6**). Sequence similarities between PfFNT and TgFNT1-3 are shown in **Table S7**. Notably, TgFNT1 is not susceptible to inhibition by MMV007839^35^, suggesting that differences at certain positions within the transport pathway may influence susceptibility and could highlight potential resistance-conferring sites in PfFNT. To explore this further and investigate whether mutation at other sites, such as V203, could destabilise inhibitor binding, we performed thermodynamic integration simulations where the sidechains of V54, V203, V216, and V220 were all mutated into a larger leucine sidechain. MMV007839 binding was destabilised in all four mutant systems, although no mutation appeared as destabilising as the V196L mutant (**Fig. 4b-c**). Lactate bound within the cavity in all systems, indicating that these mutations are unlikely to prevent PfFNT from expelling lactate from the parasite (**Fig. 4e**).

## Discussion

PfFNT is a lactate/H^⁺^ symporter that has been suggested as a potential drug target for antimalarial development due to its essential role in the disease-causing intra-erythrocytic stage of the parasite lifecycle.^11–14^ However, three mutations - G107S, V196L, and G21E - have been identified that are associated with resistance to known inhibitors such as MMV007839.^13,14,17^ Due to the narrow architecture of the PfFNT transport pathway, it has been suggested that the protein may have limited mutational flexibility within the cavity.^17^ However, the binding sites of lactate^26^ and MMV007839^19,20^ do not completely overlap, with many residues contributing to MMV007839 binding appearing to be dispensable for lactate transport (**Fig. 1c-d**). One principle of inhibitor design is the Substrate Envelope Hypothesis, which proposes that inhibitors should not make more interactions with their target proteins than the natural substrate, in order to limit the number of mutations that can be tolerated by the substrate but not the inhibitor.^36–43^ On this basis, it is likely that additional resistance-conferring mutations could occur at residues that are critical for MMV007839 binding but not for lactate transport.

In this study, we identified two additional mutations associated with resistance to MMV007839: L198M and V200L. While the previously identified G107S, V196L, and G21E mutations seem to confer resistance by sterically clashing with the inhibitor and/or altering the dynamics and free-energy landscape of the transport pathway (**Fig. 2**), the L198M and V200L mutations appear to destabilise inhibitor binding by altering the mobility of transmembrane helices that line the PfFNT cavity (**Fig. 3**). Notably, these mutations do not appear to prevent lactate transport in simulations or prevent robust growth of parasites *in vitro*. Although the frequency of lactate binding appears slightly reduced in mutant systems, it has previously been demonstrated that the G107S mutation does not cause a fitness defect despite causing ∼30-60% reduction in lactate transport.^13^ The identification of the L198M mutation demonstrates that resistance-conferring mutations in PfFNT are not restricted to residues immediately surrounding the binding site but may also occur at further away sites and destabilise inhibitor binding through subtle, allosteric changes to the transport cavity while still enabling PfFNT-mediated lactate/H^+^ transport to occur.

Notably, the L198M mutation identified in this study is located at the interface between PfFNT subunits, representing the first identified resistance-associated mutation in PfFNT that lies outside the transport pathway itself. As mutations at the oligomeric interfaces of PfFNT have not previously been explored, this finding warrants further investigation into whether additional resistance-conferring mutations may arise at these sites.

Although other interface mutations were not examined further in this study, additional potential mutations within the transport pathway were predicted using molecular docking and by selectively mutating valine residues along the transport pathway (**Fig. 4**). Notably, the V203L mutation was also predicted by molecular docking to destabilise not only MMV007839 binding, but also the binding of Compounds 7e and 8a - two recently identified PfFNT inhibitors with IC_50_ values in the nM range. Several mutations were suggested by docking to destabilise the binding of all three inhibitors (**Fig. S5**). Further study is warranted to investigate whether these mutations could arise from prolonged exposure of parasites to these inhibitors *in vitro*, and whether these mutations could give rise to resistance to one or more PfFNT inhibitors. Importantly, because our docking screen considered all possible single amino acid substitutions throughout the transport pathway, not all mutations predicted to destabilise inhibitor binding are likely to be tolerated by PfFNT. Previous studies have shown that the engineered G107V, G107C, G107A, A58V, A58L, A58F, and A224V mutations substantially impair [^14^C]-lactate uptake in PfFNT-expressing *S. cerevisiae,* reducing uptake by approximately 75-100%.^14,44^ Likewise, mutation of H230, a residue critical for lactate/H^+^ transport, to either alanine (H230A) or asparagine (H230N) abolishes lactate uptake in stop-flow assays using proteoliposomes reconstituted with PfFNT.^20^ MD simulations of PfFNT with the H230N mutation or a K35C/K177C double mutation also showed a loss of lactate binding.^26^ Consequently, some predicted resistance mutations may be evolutionarily inaccessible because of their potential detrimental effects on PfFNT function.

Cross-resistance to PfFNT inhibitors has previously been seen between the structural analogues MMV007839 and MMV000972 with the G107S mutation^13,14^, as well as MMV007839, BH267.meta, and BH296 with the G107S and V196L mutations^17^. However, it is important to note that of the five PfFNT mutations associated with resistance to MMV007839, only G107S was identified as destabilising in the docking screen, despite all five mutations destabilising MMV007839 binding in TI simulations. This finding shows that docking alone is not sufficient to accurately predict all resistance-conferring mutations, likely because it cannot capture the dynamic and allosteric effects revealed by molecular dynamics simulations. Our filtering of docking results by displacement of MMV007839 from the binding site in the wildtype cryo-EM structure also cannot account for cases where the inhibitor can likely still bind at the same site but with lowered binding affinity, as seen in MD simulations of PfFNT-G107S, PfFNT-V196L, PfFNT-G21E, PfFNT-V200L, and PfFNT-L198M (**Fig. 3d**). Nevertheless, some of the additional mutations identified by docking may still represent candidate resistance-conferring mutations worthy of further investigation. Interestingly, T106A, an engineered mutation in PfFNT that was shown from isothermal titration calorimetry experiments to reduce the binding affinity of MMV007839^19^, was identified from docking as likely to destabilise MMV007839 binding.

The two additional mutations identified in this study, together with several predicted mutations that could destabilise MMV007839 binding without impairing lactate transport, suggest that other mutations could still arise within PfFNT that only require a single base change. Notably, the discovery of the L198M mutation indicates that resistance-conferring changes may occur both within and outside the transport pathway, highlighting the need to consider potential interface mutations that could confer resistance when designing new PfFNT inhibitors.

Intriguingly, some engineered PfFNT mutations, such as K35A, F90A, F94A, and K177A have been shown to increase PfFNT-mediated lactate transport activity relative to wildtype PfFNT in stop-flow assays in reconstituted proteoliposomes.^20^ If such gain-of-function mutations were to arise in parasites, it would be interesting to investigate whether these mutations affect parasite fitness and alter the parasite’s susceptibility to PfFNT inhibitors.

The ability of parasites to tolerate at least a ∼30–60% reduction in PfFNT-mediated lactate/H^⁺^ export^13^, together with the apparent lack of growth defects associated with known resistance mutations and the rapid emergence of such mutations *in vitro* within two to six weeks^13,14,17^, suggests that PfFNT-targeting inhibitors may be particularly susceptible to the evolution of resistance if deployed clinically. Importantly, this risk is not limited to only MMV007839 and its close structural analogue MMV000972. While these two MMV compounds are toxic to mammalian cell lines^16,45^, the inhibitor BH267.meta exhibits reduced toxicity in human liver and kidney cell lines and is effective at clearing parasitemia in humanized mouse models of *P. falciparum* infection^17,45^. However, parasites carrying the PfFNT-G107S mutation are highly resistant to BH267.meta^17^, despite earlier docking predictions that this mutation would not affect inhibitor binding^19^. Furthermore, although several newly synthesised PfFNT inhibitors display improved potency over MMV007839, they all share a common pentafluoro-3-hydroxy-pent-2-en-1-one scaffold^14,15,18,45,46^, raising the likelihood that all resistance-conferring mutations identified so far could confer cross-resistance without impacting the growth of blood-stage parasites. However, it is important to consider that our assessments of parasite growth were determined from experiments performed over only one parasite intra-erythrocytic lifecycle, which would allow only major defects in parasite growth to be detected. To investigate this further, the mutant parasites could be cloned out, then their growth rates compared with that of parental parasites over several weeks in ‘competition experiments’, as previously performed with Dd2^PfFNTG107S^ parasites and their parents^13^.

Previous work by Golldack et al. has investigated whether PfFNT inhibitor design can be improved by shortening or sterically reducing the fluoroalkyl tail to evade resistance-associated mutations^14^. However, the authors demonstrated that a fluoroalkyl tail containing five fluorine atoms is optimal for inhibiting PfFNT-mediated lactate/H^⁺^ transport in PfFNT-expressing yeast, and identified a 5,5,6,6,6-pentafluoro-4-hydroxyhex-3-en-2-one pharmacophore that inhibited PfFNT-mediated [^14^C]-lactate transport in yeast with an IC_50_ of 1.9 ± 0.3 μM.^14^ Consistent with this, removal of the fluoroalkyl tail or replacement of the fluorines with other halogens reduces the potency by which the compounds inhibit lactate transport by PfFNT and/or inhibit the growth of wildtype parasites^14^, suggesting that designing potent, resistance-defying PfFNT inhibitors through modification of this moiety may be challenging.^14^

Broad anion channel inhibitors, including niflumic acid, cinnamic acid, phloretin, furosemide, α-cyano-4-hydroxy-cinnamate, α-fluorocinnamate, and 5-nitro-2-(3-phenylpropylamino)benzoic acid (NPPB) have also been shown to inhibit PfFNT-mediated lactate/H^+^ transport.^11–14,47^ Although these compounds lack specificity, they nonetheless may represent alternative scaffolds for the design of new PfFNT inhibitors.

However, all of these compounds and the more specific PfFNT inhibitors share a common problem: their large size relative to PfFNT’s substrate, lactate. Ideally, according to the Substrate Envelope Hypothesis, an inhibitor of PfFNT should be as small and similar to lactate as possible. However, as human cells also contain lactate transporters and channels^31,48,49^, making an inhibitor too similar to lactate could create challenges for selectivity and preventing off-target effects.

The selection of multiple resistance-conferring mutations in PfFNT *in vitro* highlights a potential vulnerability of PfFNT-targeting inhibitors. This does not necessarily undermine PfFNT as a viable antimalarial drug target, but efforts to do so need to be aware of this constraint. Resistance has been generated in vitro for multiple antimalarial compounds that have nevertheless found clinical utility, such as Atovaquone – an inhibitor of the cytochrome bc1 complex (Complex III) in the parasite electron transport chain^50^. The emergence and spread of resistance *in vivo* is likely to depend on additional factors, including the speed of parasite clearance, the fitness and transmissibility of resistant parasites, and pharmacokinetic and pharmacodynamic considerations. Furthermore, rational dosing strategies and the use of partner drugs may reduce the likelihood of selecting PfFNT resistance mutations in a clinical setting, or it could be inhibited by compounds that also have additional targets.

In summary, our findings demonstrate that PfFNT may have greater mutational flexibility than previously believed, with resistance-conferring mutations arising both within and outside the transport pathway (including at the interface between subunits). These mutations destabilise inhibitor binding through diverse mechanisms while not preventing lactate binding in simulations or parasite growth in *vitro*. Together, these results suggest that while PfFNT remains an essential parasite transporter and attractive potential antimalarial target, inhibitors directed against PfFNT may be inherently vulnerable to resistance if used clinically without careful consideration of evolutionary mutational constraints. More broadly, this work highlights the utility of integrating structural, biophysical, and evolutionary perspectives when evaluating membrane transport proteins as drug targets and can be used to help guide any future rational design of new PfFNT inhibitors.

## Methods

### Ethics statement

The use of human blood in this study (from anonymous donors) was approved by the Australian National University Human Research Ethics Committee (Protocol number 2017/351).

### Parasite lines and culturing

Several *P. falciparum* lines were used in this study, including the chloroquine-resistant Dd2 parental line with PfFNT-WT (Dd2^PfFNT-WT^) and a PfFNT-G107S mutant clone (Dd2^PfFNT-G107S^) generated previously from *in vitro* evolution experiments with MMV007839^13^. The remaining parasites (with V200L or L198M mutations in PfFNT) were generated in this study using the Dd2^PfFNT-WT^ parental line.

Parasites were cultured as described previously.^51,52^ Parasites were cultured with Group O or Group A (Rh+) erythrocytes, maintained at 2-4% haematocrit in a culture medium consisting of RPMI-1640 medium containing 25 mM HEPES, and supplemented with additional D-glucose (to give a final concentration of 20 mM), 200 µM hypoxanthine, 0.6% (v/v) Albumax, and 24 mg/L gentamicin. Cultures were incubated with gentle shaking at 37 °C in at atmosphere consisting of 96% N□/3% CO□/1% O□. Cultures were synchronised using D-sorbitol according to a method described previously.^53^

### Purchasing of chemicals

MMV007839 and chloroquine were purchased from MolPort and Sigma, respectively. Stock solutions were prepared in MilliQ water for chloroquine and DMSO for MMV007839, and diluted to the desired concentrations in culture medium such that the final concentration of DMSO in any experiment did not exceed 0.1% (v/v).

### MMV007839 resistance selections and pffnt sequencing

*In vitro* evolution experiments were performed using a clone of Dd2 (referred to here as Dd2^PfFNT-WT^) that was used previously to generate Dd2^PfFNT-G107S^ parasites^13^. Ten 10 mL cultures (∼4 × 10□ parasites each) were exposed to 750 nM MMV007839 (∼3 × IC□□). Cultures were checked daily for the re-emergence of parasites for the first 8 days, and the culture medium (containing 750 nM MMV007839) replaced. Thereafter, fresh medium (containing 750 nM MMV007839) and blood was supplied as needed. Cultures in which parasites ‘re-emerged’ were expanded to 50 mL, and were maintained with 750 nM MMV007839 for a total of 42 days. Cultures were discarded after 52 days if no parasites were observed.^55^

Genomic DNA was extracted from resistant parasites (isolated from their host erythrocytes using saponin as described previously^54^) and their Dd2 parent parasites using the DNeasy Plant Mini Kit (Qiagen). To sequence the entire *pffnt* gene, two PCR reactions were performed using PrimeSTAR GXL DNA polymerase. The primers used for PCR and sequencing are the same as those used previously.^13^ Sequencing was performed at the Biomolecular Resource Facility (John Curtin School of Medical Research, Australian National University).

### Parasite proliferation assays

Parasite proliferation was measured in 96-well plates containing serial dilutions of either MMV007839 or chloroquine^56^, with surviving parasites detected at the end of the assay using a DNA-intercalating dye.^55^

Assays were initiated with ring-stage *P. falciparum*-infected erythrocytes at 1% haematocrit and 0.8–1% parasitaemia. The experiments and data analysis were carried out essentially as described previously^56^, except that the duration of the assays was 72 h and chloroquine was used for the ‘zero growth’ control.

To compare the growth rates of PfFNT mutant and WT parasites in the absence of MMV007839, 10 mL cultures of infected erythrocytes (2% haematocrit, 0.8% or 1% parasitaemia) were prepared, with the parasitaemia determined again 48 h later by microscopic examination of Giemsa-stained thin blood smears.

### Analysis of experimental data

Technical replicates were averaged prior to analysis. Graphing and statistical analysis were performed using Prism 10. Statistical significance was calculated using a one-way ANOVA. An adjusted *p* value was calculated using the *post-hoc* Tukey test to assess the significance of differences between pairs of group means.

### Simulation set up

Simulation systems of PfFNT (PDB ID: 7E27)^19^ had been prepared previously.^26^ Briefly, this system contained the entire PfFNT pentamer embedded in a model POPC bilayer using CHARMM-GUI^57^, with two POPC lipids inside the central hole. Systems were solvated with 150 mM KCl, as these ions are expected to be the most abundantly found ions in the parasite cytosol (estimated concentrations of 130-149 mM^58,59^ and 48 mM^60^, respectively). The system contained ∼222,220 atoms, with overall dimensions of approximately 130 Å by 130 Å by 118 Å. Systems were visualized using the Visual Molecular Dynamics (VMD) software^61^. As the 7E27 cryo-EM structure was resolved with MMV007839 bound within the cavity, this system was chosen to study the effects of mutations on destabilizing protein-inhibitor interactions.

### General simulation methods

Topology and coordinate files for each system were generated using TLEaP with the AMBER ff19SB protein forcefield^62^, lipid21 lipid forcefield^63^, OPC water model^64^, and 12-6-4 ion parameters^65^. MMV007839 was modelled in its linear vinylogous acid tautomeric form, as resolved in the 7E27 cryo-EM structure.^19^ Parameters for MMV007839 were generated using GAFF2 and antechamber ^66^. ParmEd was used to repartition the masses of heavy atoms onto the hydrogen atoms to 3.024 Da for non-water molecules to allow a 4 fs timestep to be used.^67,68^ Subsequently, each system was prepared for MD simulations by (1) energy minimization, to eliminate steric clashes and unfavourable molecular interactions and allow the system to be brought to a local energy minimum, (2) heating, using the Langevin thermostat to slowly increase the temperature of the system to 310 K (36.85°C) over 500 ps, and (3) pressurizing, using the Monte Carlo barostat to maintain the system pressure at 1 atm over 200 ps, with the Berendsen barostat having been used for all prior preparatory phases. Throughout each of these preparatory phases, a 1 kcal/mol·Å^2^ harmonic restraint was kept on the α-carbon atoms of the protein, enabling lipids and water to accommodate the protein structure. Subsequently, restraints were incrementally released over a timeframe of 5 ns. Energy minimization and an initial heating step between -273.15 and -242.15 °C were performed using the pmemd.MPI parallel CPU implementation, while all other phases, including production, were performed using the pmemd.cuda GPU implementation (unless specified otherwise). Hydrogen bonds were constrained with SHAKE. Electrostatic interactions were calculated using the Particle Mesh Ewald (PME) summation^69^ with a real-space cut-off of 10 Å. All system preparatory phases and production simulations were performed in triplicate for each system using AMBER22.^70^

### Choice of H230 and MMV007839 protonation states

PfFNT contains several titratable residues whose pKa values have been previously assessed.^26^ Three of these residues lie near the inhibitor-binding site: D103, E229, and H230. Whereas D103 and E229 are predicted to be deprotonated at the parasite cytosolic pH (∼7.2), pKa predictions for H230 are ambiguous. Previous simulations indicate that protonation of H230 is critical for substrate transport^26^; however, it is unclear whether this is also required for MMV007839 binding.

Like H230, MMV007839 can also adopt multiple protonation states (**Fig. 1e**). In its linear form the compound contains two protonatable oxygen atoms, one on the aromatic ring (“ring oxygen”) and one near the fluoroalkyl tail (“tail oxygen”), giving four possible protonation states: both protonated, ring-only, tail-only, or neither protonated. Initial attempts to predict the pK_a_ of the titratable groups of MMV007839 with MolGpKa and Marvinsketch suggested the tail oxygen is likely deprotonated (**Fig. S7a**). Notably, this group on MMV007839 is within hydrogen bonding distance of a hydroxyl group on T106, supporting the oxygen near the fluoroalkyl tail being deprotonated to participate in a hydrogen bond (**Fig. S7b**). To determine which protonation state combination leads to the most favourable interactions between the protein and the inhibitor, we performed initial molecular docking and molecular dynamics simulations for all possible combinations of H230 and MMV007839 protonation state. The starting structures for each protonation state were generated by protonating all possible sites on H230 and MMV007839 and manually removing protons where required to yield the other protonation state combinations.

Molecular docking was performed using AutoDock Vina v1.2.5.^71,72^ Protein and ligand files were converted to pdbqt format using AutoDockTools^73^ and OpenBabel^74^, respectively. The search space used in all docking runs was limited to a 25 x 25 x 30 Å region, determined using AutoDockTools, where the centre of the box was adjusted to encompass the intracellular side of the transport pathway within a single subunit. Initial docking for protonation state tests were performed within five pseudoreplicates by docking to each subunit with an exhaustiveness of 200. The top ranked poses were used for analysing the most likely protonation state combinations. RMSD of docked poses relative to the position of MMV007839 within the 7E27 cryo-EM structure were measured using CPPTRAJ^75^ (**Fig. S7c**).

MD simulations were performed using a modified version of the simulation protocol described above. Following the standard heating steps, the system was further heated from 310 K to 330 K in 2 K increments every 2 ns and simulations were subsequently performed for 100 ns at 330 K to facilitate escape of MMV007839 from local energy minima. The RMSD of MMV007839 and its distance from the extracellular constriction site (formed by H230, F90, and F223) were measured using CPPTRAJ (**Fig. S7d**). In addition, the χ1 (N–CA–CB–CG) and χ2 (CA–CB–CG–ND1) side-chain dihedral angles of H230 were analysed to determine which protonation state best preserved the conformation observed in the cryo-EM structure (**Fig. S7e**). These analyses supported the use of a H230 protonated at the ε-nitrogen and an MMV007839 molecule protonated at the ring oxygen for subsequent simulations. Unless specified otherwise, these protonation states were used for all subsequent simulations of PfFNT with MMV007839 present.

### Computational generation of mutant PfFNT structures

Selected residues in the WT 7E27 structure were mutated in all five PfFNT subunits using the ‘Mutate Model’ function within the MODELLER program^76^, which iteratively builds in the new mutant side chains through a series of energy minimisation steps. Residue protonation states were then re-assigned using TLEaP.

### Simulations with MMV007839 bound within the cavity

PfFNT-WT simulation systems were mutated using MODELLER to introduce select mutations (G107S, G21E, V196L, V200L, and L198M) into all five PfFNT subunits. Simulations of systems were performed as described above in triplicate for each system for 1 μs per replicate. The RMSD of MMV007839 within the cavity over time and the distance between the COM of MMV007839 and the COM of the extracellular constriction were measured using CPPTRAJ (**Fig. S1a-b**).

### MM/GBSA calculations

MM/GBSA calculations were performed using the mmgbsa.py script from AmberTools2025. For each replicate from the PfFNT-WT, PfFNT-G107S, PfFNT-G21E, PfFNT-V196L, PfFNT-V200L, and PfFNT-L198M simulation systems with MMV007839 bound within the cavity, cpptraj was used to additionally prepare topologies and trajectories containing only the protein, only MMV007839, and only the protein and MMV007839 from the final 200 ns of each replicate. MM/GBSA was run using the Generalised Born model with igb = 5 and a salt concentration of 0.150 M. Per-residue energy decomposition was performed with idecomp = 1. The entropic contributions were not included in the MM/GBSA calculations. For each system, average per-residue decomposition values were compared against the corresponding average WT values to estimate the change in residue-wise binding contribution, calculated as ΔΔG = ΔG mutant – ΔG WT. The data from the MM/GBSA calculations are shown in **Fig. S1c**.

### Steered MD and Umbrella sampling simulations for PfFNT-WT and PfFNT-G107S

To generate starting coordinates for umbrella sampling (US), steered MD (SMD) simulations were performed for the PfFNT-WT and PfFNT-G107S systems using a protocol described previously.^26^ For each system, the final frame of one replicate of the simulations with MMV007839 bound within the cavity in each subunit was used as a starting frame for SMD. For both SMD and US, the collective variable used for the intracellular side of the protein was the distance between the COM of the backbone atoms of extracellular constriction and the COM of MMV007839.

For the SMD, a force constant of 25 kcal/mol·Å^2^ was used to move MMV007839 in 0.5 Å increments every 2 ns in all five subunits simultaneously. AMBER restart files were written every 2 ns of simulation to obtain starting coordinates for US along the pathway from the cavity to bulk. For both systems, one-dimensional umbrella sampling (US) was used to sample the region between the cavity and the bulk solution on the intracellular side. A total of 20 windows were used, spaced 0.5 Å apart – covering the region from the transport cavity to the bulk solution on the intracellular side. All umbrella sampling (US) trajectories were unbiassed, and the potential of mean force (PMF) was computed using a variant of the Weighted Histogram Analysis Method (WHAM) as implemented by Alan Grossfield.^77^ WHAM was run under non-periodic conditions using 500 bins and a convergence criterion of 1 × 10^⁻^. The histogram limits were defined by the minimum and maximum values of the inhibitor’s position along the reaction coordinate. To estimate the appropriate equilibration period to exclude, PMFs were recalculated for each subunit after progressively trimming data from the beginning of the trajectories (**Figure S8**). After determining equilibration time, PMFs were recalculated again using increasingly more simulation time to assess convergence (**Figure S9**). Each window was simulated for 100 ns with a force constant of 15 kcal/mol·Å^2^. Data from 50-100 ns was used for generating the final PMF curves.

### Thermodynamic integration simulations

Thermodynamic integration (TI) was used to predict the relative binding free energy (ΔΔG) of MMV007839 for seven PfFNT mutants compared with WT. ΔΔG was calculated from the thermodynamic cycle as ΔG□ − ΔG□, where ΔG□ is the free energy change for the WT→mutant transformation in the ligand-bound state and ΔG□ is the corresponding change in the apo state. TI simulations were performed in pmemd.cuda (AMBER22) using dual-topology models generated in pdb4amber.

Dual-topology parameter and restart files were generated in TLEaP using ff14SB for protein^78^, LIPID21 for lipids^63^, TIP3P water^79^, 12-6-4 ion parameters^65^, and previously derived MMV007839 parameters. Non-mutated residues in the selected subunit were merged using ParmED TIMerge, leaving only the mutated residue under λ-control with soft-core potentials applied.

Twenty-three λ windows were used to cover the space between λ = 0 and λ = 1 in steps of 0.05, with additional windows at λ = 0.96 and λ = 0.98. Each window was simulated triplicate for both ΔG□ and ΔG□. Windows were simulated for 50 ns each for the PfFNT-G107S, PfFNT-V196L, PfFNT-G21E, PfFNT-L198M, and PfFNT-V200L systems, or for 25 ns each for all other systems. Simulations were performed at 310 K and 1 bar using Langevin dynamics with a collision frequency of 5 ps⁻¹ and Monte Carlo barostat coupling. Long-range electrostatics were treated with PME with a 10 Å real-space cutoff. A 1 fs timestep was used without SHAKE constraints on bonds involving hydrogen. Convergence was assessed by the stability of the cumulative dV/dλ values and by calculating whether ΔΔG stabilised with the inclusion of increasing amounts of simulation time (**Fig. S10**). Free energies were obtained by trapezoidal integration of dV/dλ across λ using alchemlyb^80^, and ΔΔG values were calculated as ΔG□ − ΔG□.

### Molecular docking

Computational screening of MMV007839 binding poses to different mutant structures of PfFNT was performed using AutoDock Vina v1.2.5 with a similar docking search space to what was described above for the protonation state docking^71,72^. Docking was performed using flexible docking^71^, where the inhibitor and the sidechains of the intracellular constriction as well as of the mutant residue in each docking pose were allowed to be flexible. Docking to each mutant structure was performed with five independent replicates with an exhaustiveness value of 400.

Each independent docking run produced 9 different poses. Poses from all five docking replicates were combined and clustered using K-means clustering within CPPTRAJ. A representative pose from the top ranked cluster with the minimum distance from the centroid of that cluster was used for further analysis, unless specified otherwise. CPPTRAJ was used to calculate the RMSD between the top cluster docked pose and the pose of MMV007839 in the 7E27 cryo-EM structure, as well as the distance between the COM of the docked pose and the extracellular constrictions.

Docking results were filtered for mutations that required only a single base change. Mutations were considered destabilising if the RMSD of the docked pose relative to the position of MMV007839 in the 7E27 cryo-EM structure was greater than 3.5 Å.

The same docking protocol was used for running the docking pipeline with compounds 7e and 8a. To measure the RMSD of the docked poses of these compounds relative to the position of MMV007839 within the cryo-EM binding site, RDKit^81^ was used to calculate RMSD based on the maximum common substructure between each compound and MMV007839.

### Analysis of computational data

Unless stated otherwise, analyses were conducted in Python using MDAnalysis^82^, with NumPy^83^ and Pandas^84^ for data processing. Figures were generated using Matplotlib^85^ and Seaborn^86^. Statistical analyses were performed in RStudio (R version 4.4.1). Binding events were analysed using a binomial generalised linear model (GLM) with a logit link function, with mutation treated as a fixed effect. Pairwise comparisons against PfFNT-WT were performed using Dunnett-adjusted post hoc tests implemented in the emmeans package. Model diagnostics were assessed using the DHARMa package.^87–89^

## Author Contribution Statements

Parasite culturing: A.P.

*In vitro* evolution experiments: A.P.

Parasite proliferation and fitness assays: A.P.

Docking: A.P., C.W.

Flooding simulations: C.W.

Thermodynamic integration simulations: C.W., A.P.

Umbrella sampling simulations: C.W.

All other MD simulations: A.P., C.W.

Figure making: C.W., A.P.

Writing and editing: C.W., A.P., A.M.L., B.C.

Funding acquisition: A.M.L, B.C.

Supervision: C.W., A.M.L, B.C.

Conceptualisation: C.W., A.P., A.M.L., B.C.

## Acknowledgements

We would like to acknowledge Deyun Qiu for assistance with experiments and sequencing, and Jonathan Rocco for assistance with statistical analysis. We would also like to thank John Tanner for useful discussions on the substrate envelope hypothesis and docking. This work was supported by resources and services provided by the National Computational Infrastructure (NCI) and the Pawsey Supercomputing Research Centre, funded by the Australian Government (project g15). A.P. acknowledges a Medical Advances Without Animals (MAWA) Trust Honours Scholarship. C.W. acknowledges an Australian Government Research Training Program PhD Scholarship. A.M.L. acknowledges funding from Australian National Health and Medical Research Council (GNT2028714). B.C. acknowledges funding from Australian Research Council (DP250100893). We wish to thank the Canberra Branch of the Australian Red Cross Lifeblood for the provision of blood, and the ANU Biomolecular Resource Facility for sequencing.

## References

1. World Health Organisation. World malaria report 2025. https://www.who.int/publications/i/item/9789240117822 (2025).

2. Lalremruata, A. et al. Species and genotype diversity of Plasmodium in malaria patients from Gabon analysed by next generation sequencing. Malar. J. 16, 398 (2017).

3. Oladipo, H. J. et al. Increasing challenges of malaria control in sub-Saharan Africa: Priorities for public health research and policymakers. Ann. Med. Surg. (Lond). 81, (2022).

4. Ishengoma, D. S. et al. Urgent action is needed to confront artemisinin partial resistance in African malaria parasites. Nat. Med. 30, 1807–1808 (2024).

5. Balikagala, B. et al. Evidence of Artemisinin-Resistant Malaria in Africa. New England Journal of Medicine 385, 1163–1171 (2021).

6. van der Pluijm, R. W. et al. Triple artemisinin-based combination therapies versus artemisinin-based combination therapies for uncomplicated Plasmodium falciparum malaria: a multicentre, open-label, randomised clinical trial. The Lancet 395, 1345–1360 (2020).

7. van der Pluijm, R. W., Amaratunga, C., Dhorda, M. & Dondorp, A. M. Triple Artemisinin-Based Combination Therapies for Malaria – A New Paradigm? Trends in Parasitology vol. 37 15–24 Preprint at 10.1016/j.pt.2020.09.011 (2021).

8. Zhu, L. et al. Artemisinin resistance in the malaria parasite, Plasmodium falciparum, originates from its initial transcriptional response. Commun. Biol. 5, 1–13 (2022).

9. Phyo, A. P. et al. Declining Efficacy of Artemisinin Combination Therapy Against P. Falciparum Malaria on the Thai-Myanmar Border (2003-2013): The Role of Parasite Genetic Factors. Clinical Infectious Diseases 63, 784–791 (2016).

10. Siqueira-Neto, J. L. et al. Antimalarial drug discovery: progress and approaches. Nature Reviews Drug Discovery 2023 22:10 22, 807–826 (2023).

11. Marchetti, R. V. et al. A lactate and formate transporter in the intraerythrocytic malaria parasite, Plasmodium falciparum. Nature Communications 2015 6:1 6, 1–7 (2015).

12. Wu, B. et al. Identity of a Plasmodium lactate/H+ symporter structurally unrelated to human transporters. Nature Communications 2015 6:1 6, 1–8 (2015).

13. Hapuarachchi, S. V. et al. The Malaria Parasite’s Lactate Transporter PfFNT Is the Target of Antiplasmodial Compounds Identified in Whole Cell Phenotypic Screens. PLoS Pathog. 13, 1–24 (2017).

14. Golldack, A. et al. Substrate-analogous inhibitors exert antimalarial action by targeting the Plasmodium lactate transporter PfFNT at nanomolar scale. PLoS Pathog. 13, 1–18 (2017).

15. Walloch, P., Hansen, C., Priegann, T., Schade, D. & Beitz, E. Pentafluoro-3-hydroxy-pent-2-en-1-ones Potently Inhibit FNT-Type Lactate Transporters from all Five Human-Pathogenic Plasmodium Species. https://doi.org/10.1002/cmdc.202000952 (2020) doi:10.1002/cmdc.202000952.

16. Van Voorhis, W. C. et al. Open Source Drug Discovery with the Malaria Box Compound Collection for Neglected Diseases and Beyond. PLoS Pathog. 12, e1005763 (2016).

17. Davies, H. et al. The Plasmodium Lactate/H+ Transporter PfFNT Is Essential and Druggable In Vivo. Antimicrob. Agents Chemother. https://doi.org/10.1128/AAC.00356-23 (2023) doi:10.1128/AAC.00356-23.

18. Nerlich, C., Epalle, N. H., Seick, P. & Beitz, E. Discovery and development of inhibitors of the plasmodial fnt-type lactate transporter as novel antimalarials. Pharmaceuticals 14, 1191 (2021).

19. Peng, X. et al. Structural characterization of the Plasmodium falciparum lactate transporter PfFNT alone and in complex with antimalarial compound MMV007839 reveals its inhibition mechanism. PLoS Biol. 19, 1–19 (2021).

20. Lyu, M., Su, C., Kazura, J. W. & Yu, E. W. Structural basis of transport and inhibition of the Plasmodium falciparum transporter PfFNT. EMBO Rep. 22, 1–12 (2021).

21. Waight, A. B., Love, J. & Wang, D.-N. N. Structure and mechanism of a pentameric formate channel. Nat. Struct. Mol. Biol. 17, 31–38 (2010).

22. Lü, W. et al. pH-dependent gating in a FocA formate channel. Science (1979). 332, 352–354 (2011).

23. Lü, W. et al. The formate channel FocA exports the products of mixed-acid fermentation. Proc. Natl. Acad. Sci. U. S. A. 109, 13254–13259 (2012).

24. Wang, Y. et al. Structure of the formate transporter FocA reveals a pentameric aquaporin-like channel. Nature 462, 467–472 (2009).

25. Czyzewski, B. K. & Wang, D. N. Identification and characterization of a bacterial hydrosulphide ion channel. Nature 2012 483:7390 483, 494–497 (2012).

26. Wallis, C. et al. A proton transfer mechanism in the malaria parasite lactate/H+ symporter reveals a channel-like transporter without conformational changes. bioRxiv 2026.02.05.704099 (2026) doi:10.64898/2026.02.05.704099.

27. Deng, D. et al. Crystal structure of the human glucose transporter GLUT1. Nature 2014 510:7503 510, 121–125 (2014).

28. Woodrow, C. J., Penny, J. I. & Krishna, S. Intraerythrocytic Plasmodium falciparum expresses a high affinity facilitative hexose transporter. Journal of Biological Chemistry 274, 7272–7277 (1999).

29. Krishna, S., Woodrow, C. J., Burchmore, R. J. S., Saliba, K. J. & Kirk, K. Hexose Transport in Asexual Stages of Plasmodium falciparum and Kinetoplastidae. Parasitology Today 16, 516–521 (2000).

30. Dunn, C. R. et al. The structure of lactate dehydrogenase from Plasmodium falciparum reveals a new target for anti-malarial design. Nat. Struct. Biol. 3, 912–915 (1996).

31. Poole, R. C. & Halestrap, A. P. Transport of lactate and other monocarboxylates across mammalian plasma membranes. 10.1152/ajpcell.1993.264.4.C761 264, (1993).

32. Ginsburg, H., Krugliak, M., Eidelman, O. & Ioav Cabantchik, Z. New permeability pathways induced in membranes of Plasmodium falciparum infected erythrocytes. Mol. Biochem. Parasitol. 8, 177–190 (1983).

33. Krugliak, M. & Ginsburg, H. The evolution of the new permeability pathways in Plasmodium falciparum—infected erythrocytes—a kinetic analysis. Exp. Parasitol. 114, 253–258 (2006).

34. Nerlich, C., et al. Addressing the Intracellular Vestibule of the Plasmodial Lactate Transporter PfFNT by p-Substituted Inhibitors Amplifies In Vitro Activity. J. Med. Chem. 67, 18368–18383 (2024).

35. Zeng, J. M. et al. Identifying the major lactate transporter of Toxoplasma gondii tachyzoites. Sci. Rep. 11, 1–11 (2021).

36. Shaqra, A. M. et al. Defining the substrate envelope of SARS-CoV-2 main protease to predict and avoid drug resistance. Nature Communications 2022 13:1 13, 3556- (2022).

37. Shen, Y. et al. Testing the Substrate-Envelope Hypothesis with Designed Pairs of Compounds. ACS Chem. Biol. 8, 2433–2441 (2013).

38. Romano, K. P. et al. Molecular mechanisms of viral and host cell substrate recognition by hepatitis C virus NS3/4A protease. J. Virol. 85, 6106–6116 (2011).

39. Romano, K. P., Ali, A., Royer, W. E. & Schiffer, C. A. Drug resistance against HCV NS3/4A inhibitors is defined by the balance of substrate recognition versus inhibitor binding. Proc. Natl. Acad. Sci. U. S. A. 107, 20986–20991 (2010).

40. Kairys, V., Gilson, M. K., Lather, V., Schiffer, C. A. & Fernandes, M. X. Toward the design of mutation-resistant enzyme inhibitors: further evaluation of the substrate envelope hypothesis. Chem. Biol. Drug Des. 74, 234–245 (2009).

41. Nalam, M. N. L. et al. Evaluating the substrate-envelope hypothesis: structural analysis of novel HIV-1 protease inhibitors designed to be robust against drug resistance. J. Virol. 84, 5368–5378 (2010).

42. Chellappan, S., Kairys, V., Fernandes, M. X., Schiffer, C. & Gilson, M. K. Evaluation of the substrate envelope hypothesis for inhibitors of HIV-1 protease. Proteins 68, 561–567 (2007).

43. King, N. M., Prabu-Jeyabalan, M., Nalivaika, E. A. & Schiffer, C. A. Combating susceptibility to drug resistance: Lessons from HIV-1 protease. Chem. Biol. 11, 1333–1338 (2004).

44. Wiechert, M., Erler, H., Golldack, A. & Beitz, E. A widened substrate selectivity filter of eukaryotic formate-nitrite transporters enables high-level lactate conductance. FEBS Journal 284, 2663–2673 (2017).

45. Nerlich, C., et al. Addressing the Intracellular Vestibule of the Plasmodial Lactate Transporter PfFNT by p-Substituted Inhibitors Amplifies In Vitro Activity. J. Med. Chem. 67, 18368–18383 (2024).

46. Walloch, P. et al. Introduction of Scaffold Nitrogen Atoms Renders Inhibitors of the Malarial l -Lactate Transporter, PfFNT, Effective against the Gly107Ser Resistance Mutation. J. Med. Chem. 63, 9731–9741 (2020).

47. Elliott, J. L., Saliba, K. J. & Kirk, K. Transport of lactate and pyruvate in the intraerythrocytic malaria parasite, Plasmodium falciparum. Biochemical Journal 355, 733 (2001).

48. Geistlinger, K., Schmidt, J. D. R. & Beitz, E. Lactic Acid Permeability of Aquaporin-9 Enables Cytoplasmic Lactate Accumulation via an Ion Trap. . 12, (2022).

49. Wang, N. et al. Structural basis of human monocarboxylate transporter 1 inhibition by anti-cancer drug candidates. Cell 184, 370–383.e13 (2021).

50. Fry, M. & Pudney, M. Site of action of the antimalarial hydroxynaphthoquinone, 2-[trans-4-(4’-chlorophenyl) cyclohexyl]-3- hydroxy-1,4-naphthoquinone (566C80). Biochem. Pharmacol. 43, 1545–1553 (1992).

51. Allen, R. J. W. & Kirk, K. Plasmodium falciparum culture: The benefits of shaking. Mol. Biochem. Parasitol. 169, 63–65 (2010).

52. Trager, W. & Jensen, J. B. Human Malaria Parasites in Continuous Culture. Science (1979). 193, 673–675 (1976).

53. Lambros, C. & Vanderberg, J. P. Synchronization of Plasmodium falciparum erythrocytic stages in culture. Journal of Parasitology 65, 418–420 (1979).

54. Saliba, K. J., Horner, H. A. & Kirk, K. Transport and metabolism of the essential vitamin pantothenic acid in human erythrocytes infected with the malaria parasite Plasmodium falciparum. Journal of Biological Chemistry 273, 10190–10195 (1998).

55. Smilkstein, M., Sriwilaijaroen, N., Kelly, J. X., Wilairat, P. & Riscoe, M. Simple and inexpensive fluorescence-based technique for high-throughput antimalarial drug screening. Antimicrob. Agents Chemother. 48, 1803–1806 (2004).

56. Spry, C. et al. Pantothenamides Are Potent, On-Target Inhibitors of Plasmodium falciparum Growth When Serum Pantetheinase Is Inactivated. PLoS One 8, e54974 (2013).

57. Lee, J. et al. CHARMM-GUI Input Generator for NAMD, GROMACS, AMBER, OpenMM, and CHARMM/OpenMM Simulations Using the CHARMM36 Additive Force Field. J. Chem. Theory Comput. 12, 405–413 (2016).

58. Winterberg, M. & Kirk, K. A high-sensitivity HPLC assay for measuring intracellular Na+ and K+ and its application to Plasmodium falciparum infected erythrocytes. Scientific Reports 2016 6:1 6, 1–6 (2016).

59. Mauritz, J. M. A. et al. X-Ray Microanalysis Investigation of the Changes in Na, K, and Hemoglobin Concentration in Plasmodium falciparum-Infected Red Blood Cells. Biophys. J. 100, 1438 (2011).

60. Henry, R. I. et al. An acid-loading chloride transport pathway in the intraerythrocytic malaria parasite, Plasmodium falciparum. Journal of Biological Chemistry 285, 18615–18626 (2010).

61. Humphrey, W., Dalke, A. & Schulten, K. VMD: visual molecular dynamics. J. Mol. Graph. 14, 33–38 (1996).

62. Tian, C. et al. Ff19SB: Amino-Acid-Specific Protein Backbone Parameters Trained against Quantum Mechanics Energy Surfaces in Solution. J. Chem. Theory Comput. 16, 528–552 (2020).

63. Dickson, C. J., Walker, R. C. & Gould, I. R. Lipid21: Complex Lipid Membrane Simulations with AMBER. J. Chem. Theory Comput. 18, 1726–1736 (2022).

64. Izadi, S., Anandakrishnan, R. & Onufriev, A. V. Building Water Models: A Different Approach. Journal of Physical Chemistry Letters 5, 3863–3871 (2014).

65. Li, P. Bridging the 12-6-4 Model and the Fluctuating Charge Model. Front. Chem. 9, 551 (2021).

66. Wang, J., Wolf, R. M., Caldwell, J. W., Kollman, P. A. & Case, D. A. Development and testing of a general amber force field. J. Comput. Chem. 25, 1157–1174 (2004).

67. Balusek, C. et al. Accelerating Membrane Simulations with Hydrogen Mass Repartitioning. J. Chem. Theory Comput. 15, 4673–4686 (2019).

68. Hopkins, C. W., Le Grand, S., Walker, R. C. & Roitberg, A. E. Long-time-step molecular dynamics through hydrogen mass repartitioning. J. Chem. Theory Comput. 11, 1864–1874 (2015).

69. Essmann, U. et al. A smooth particle mesh Ewald method. J. Chem. Phys. 103, 8577 (1998).

70. Case, D. A. et al. Amber 2022 Reference Manual ˃ SPbU Researchers Portal. Preprint at https://pureportal.spbu.ru/en/publications/amber-2022-reference-manual(ee4bb84a-9d73-44c5-bde3-76d3955ae7f1)/export.html (2022).

71. Eberhardt, J., Santos-Martins, D., Tillack, A. F. & Forli, S. AutoDock Vina 1.2.0: New Docking Methods, Expanded Force Field, and Python Bindings. J. Chem. Inf. Model. 61, 3891–3898 (2021).

72. Trott, O. & Olson, A. J. AutoDock Vina: Improving the speed and accuracy of docking with a new scoring function, efficient optimization, and multithreading. J. Comput. Chem. 31, NA-NA (2009).

73. Morris, G. M. et al. AutoDock4 and AutoDockTools4: Automated Docking with Selective Receptor Flexibility. J. Comput. Chem. 30, 2785 (2009).

74. O’Boyle, N. M. et al. Open Babel: An Open chemical toolbox. J. Cheminform. 3, 1–14 (2011).

75. Roe, D. R. & Cheatham, T. E. PTRAJ and CPPTRAJ: Software for Processing and Analysis of Molecular Dynamics Trajectory Data. J. Chem. Theory Comput. https://doi.org/10.1021/ct400341p (2013) doi:10.1021/ct400341p.

76. Eswar, N. et al. Comparative Protein Structure Modeling Using Modeller. Current protocols in bioinformatics / editoral board, Andreas D. Baxevanis … [et al.] 0 5, Unit (2006).

77. Grossfield, A. An implementation of WHAM: the Weighted Histogram Analysis Method Version 2.0.11. http://membrane. (2018).

78. Maier, J. A. et al. ff14SB: Improving the Accuracy of Protein Side Chain and Backbone Parameters from ff99SB. J. Chem. Theory Comput. 11, 3696–3713 (2015).

79. Jorgensen, W. L., Chandrasekhar, J., Madura, J. D., Impey, R. W. & Klein, M. L. Comparison of simple potential functions for simulating liquid water. J. Chem. Phys. 79, 926–935 (1983).

80. Wu, Z. et al. alchemlyb: the simple alchemistry library. J. Open Source Softw. 9, 6934 (2024).

81. Landrum, G. RDKit: Open-Source Cheminformatics Software. Preprint at (2016).

82. Michaud-Agrawal, N., Denning, E. J., Woolf, T. B. & Beckstein, O. MDAnalysis: A toolkit for the analysis of molecular dynamics simulations. J. Comput. Chem. 32, 2319–2327 (2011).

83. Harris, C. R. et al. Array programming with NumPy. Nature 2020 585:7825 585, 357–362 (2020).

84. Mckinney, W. Data Structures for Statistical Computing in Python. (2010).

85. Hunter, J. Matplotlib: A 2D Graphics Environment. Comput. Sci. Eng. 9, (2007).

86. Waskom, M. L. seaborn: statistical data visualization. J. Open Source Softw. 6, 3021 (2021).

87. Hartig, F. DHARMa: Residual Diagnostics for Hierarchical (Multi-Level / Mixed) Regression Models. Preprint at (2026).

88. Lenth, R. V. & Piaskowski, J. emmeans: Estimated Marginal Means, aka Least-Squares Means. Preprint at (2026).

89. RStudio Team. RStudio: Integrated Development Environment for R. Posit Software. Preprint at (2026).

