## Supplementary Information for "Mutations within and outside the PfFNT transport pathway drive MMV007839 resistance in *Plasmodium falciparum*"

**Table S1. IC_50_ values for MMV007839 calculated from the parasite proliferation assays for which dose-response curves are shown in Fig. 3a.**

| **Parasite culture** | **IC_50_ (µM)** | | | |
| --- | --- | --- | --- | --- |
|  | **Replicate 1** | **Replicate 2** | **Replicate 3** | **Mean +/- SEM** |
| Parent (Dd2^PfFNT-WT^) | 0.27 | 0.26 | 0.31 | 0.28 ± 0.02 |
| Dd2^PfFNT-G107S^ | 72 | 66 | 57 | 65 ± 5 |
| Dd2^PfFNT-L198M^ | 1.73 | 1.72 | 1.88 | 1.78 ± 0.05 |
| Dd2^PfFNT-V200L^ | 3.48 | 3.69 | 3.95 | 3.71 ± 0.14 |

**Table S2. IC_50_ values for chloroquine calculated from the parasite proliferation assays for which dose-response curves are shown in Fig. S2d.**

| **Parasite culture** | **IC_50_ (nM)** | | | |
| --- | --- | --- | --- | --- |
|  | **Replicate 1** | **Replicate 2** | **Replicate 3** | **Mean +/- SEM** |
| Parent (Dd2^PfFNT-WT^) | 98 | 112 | 147 | 119 ± 15 |
| Dd2^PfFNT-G107S^ | 106 | 103 | 139 | 116 ± 11 |
| Dd2^PfFNT-L198M^ | 98 | 129 | 134 | 120 ± 11 |
| Dd2^PfFNT-V200L^ | 92 | 125 | 147 | 116 ± 12 |

**Table S3. Parasitaemia (%) of each parasite line from fitness assays.**

| **Parasite culture** | **Replicate 1** | **Replicate 2** | **Replicate 3** | **Mean +/- SEM** |
| --- | --- | --- | --- | --- |
| Parent (Dd2^PfFNT-WT^) | 7.8 | 7.7 | 10.7 | 8.7 ± 1.0 |
| Dd2^PfFNT-G107S^ | 7.3 | 9.3 | 8.9 | 8.5 ± 0.6 |
| Dd2^PfFNT-L198M^ | 7.5 | 9.6 | 11.3 | 9.5 ± 1.1 |
| Dd2^PfFNT-V200L^ | 8.7 | 10.5 | 9.5 | 9.6 ± 0.5 |

**Table S4. ΔΔG values (kJ/mol) from TI calculations.**

| **System** | **Replicate 1** | **Replicate 2** | **Replicate 3** | **Mean** ± **SEM** |
| --- | --- | --- | --- | --- |
| PfFNT-G107S | 18.6 | 16.5 | 18.8 | 18.0 ± 0.7 |
| PfFNT-V196L | 16.8 | 20.5 | 17.5 | 18.3 ± 1.1 |
| PfFNT-G21E | 9.1 | 12.6 | 9.5 | 10.4 ± 1.1 |
| PfFNT-L198M | 3.9 | 7.0 | 5.2 | 5.3 ± 0.9 |
| PfFNT-V200L | 6.0 | 4.5 | 6.8 | 5.7 ± 0.7 |
| PfFNT-V54L | 8.0 | 7.9 | 8.5 | 8.1 ± 0.2 |
| PfFNT-V203L | 11.3 | 12.6 | 16.4 | 13.4 ± 1.5 |
| PfFNT-V216L | 16.0 | 14.9 | 15.6 | 18.3 ± 1.1 |
| PfFNT-V220L | 9.7 | 7.6 | 9.0 | 8.8 ± 0.6 |

**Table S5. Lactate binding events from mutant flooding simulations.** Binding events from PfFNT-WT flooding simulations are from Wallis et al.^1^

| **Simulation System** | **Replicate** | **Binding Events** | **Mean** | **SEM** |
| --- | --- | --- | --- | --- |
| PfFNT-WT | 1 | 3 | 2.8 | 0.2 |
| PfFNT-WT | 2 | 3 |  |  |
| PfFNT-WT | 3 | 2 |  |  |
| PfFNT-WT | 4 | 3 |  |  |
| PfFNT-WT | 5 | 3 |  |  |
| PfFNT-WT | 6 | 2 |  |  |
| PfFNT-WT | 7 | 3 |  |  |
| PfFNT-WT | 8 | 3 |  |  |
| PfFNT-G107S | 1 | 1 | 0.7 | 0.3 |
| PfFNT-G107S | 2 | 1 |  |  |
| PfFNT-G107S | 3 | 0 |  |  |
| PfFNT-V196L | 1 | 0 | 1.0 | 0.6 |
| PfFNT-V196L | 2 | 1 |  |  |
| PfFNT-V196L | 3 | 2 |  |  |
| PfFNT-G21E | 1 | 0 | 0.7 | 0.3 |
| PfFNT-G21E | 2 | 1 |  |  |
| PfFNT-G21E | 3 | 1 |  |  |
| PfFNT-L198M | 1 | 1 | 1.3 | 0.3 |
| PfFNT-L198M | 2 | 1 |  |  |
| PfFNT-L198M | 3 | 2 |  |  |
| PfFNT-V200L | 1 | 0 | 0.7 | 0.3 |
| PfFNT-V200L | 2 | 1 |  |  |
| PfFNT-V200L | 3 | 1 |  |  |
| PfFNT-V54L | 1 | 3 | 2.3 | 0.7 |
| PfFNT-V54L | 2 | 3 |  |  |
| PfFNT-V54L | 3 | 1 |  |  |
| PfFNT-V203L | 1 | 1 | 1.0 | 0.0 |
| PfFNT-V203L | 2 | 1 |  |  |
| PfFNT-V203L | 3 | 1 |  |  |
| PfFNT-V216L | 1 | 1 | 1.7 | 0.7 |
| PfFNT-V216L | 2 | 3 |  |  |
| PfFNT-V216L | 3 | 1 |  |  |
| PfFNT-V220L | 1 | 2 | 1.0 | 0.6 |
| PfFNT-V220L | 2 | 0 |  |  |
| PfFNT-V220L | 3 | 1 |  |  |

**Table S6. Adjusted p-values for comparisons of mutant flooding systems relative to wildtype.** Statistical significance was assessed using a binomial GLM with a logit link function, with mutation treated as a fixed effect, followed by Dunnett-adjusted post hoc comparisons against WT.

| **Simulation System** | ***p* value** |
| --- | --- |
| PfFNT-G107S | 0.0822 |
| PfFNT-G21E | 0.0822 |
| PfFNT-L198M | 0.3528 |
| PfFNT-V196L | 0.1708 |
| PfFNT-V200L | 0.0822 |
| PfFNT-V203L | 0.1708 |
| PfFNT-V216L | 0.6152 |
| PfFNT-V220L | 0.1708 |
| PfFNT-V54L | 0.9793 |

**Table S7. Sequence similarity between PfFNT and TgFNT1-3.** Percent sequence identity was calculated using the pairwise alignment tool in JalView.^2^

| **Protein Comparison** | **Percent Sequence Identity** |
| --- | --- |
| PfFNT – TgFNT1 | 32.54 |
| PfFNT – TgFNT2 | 33.76 |
| PfFNT – TgFNT3 | 36.00 |

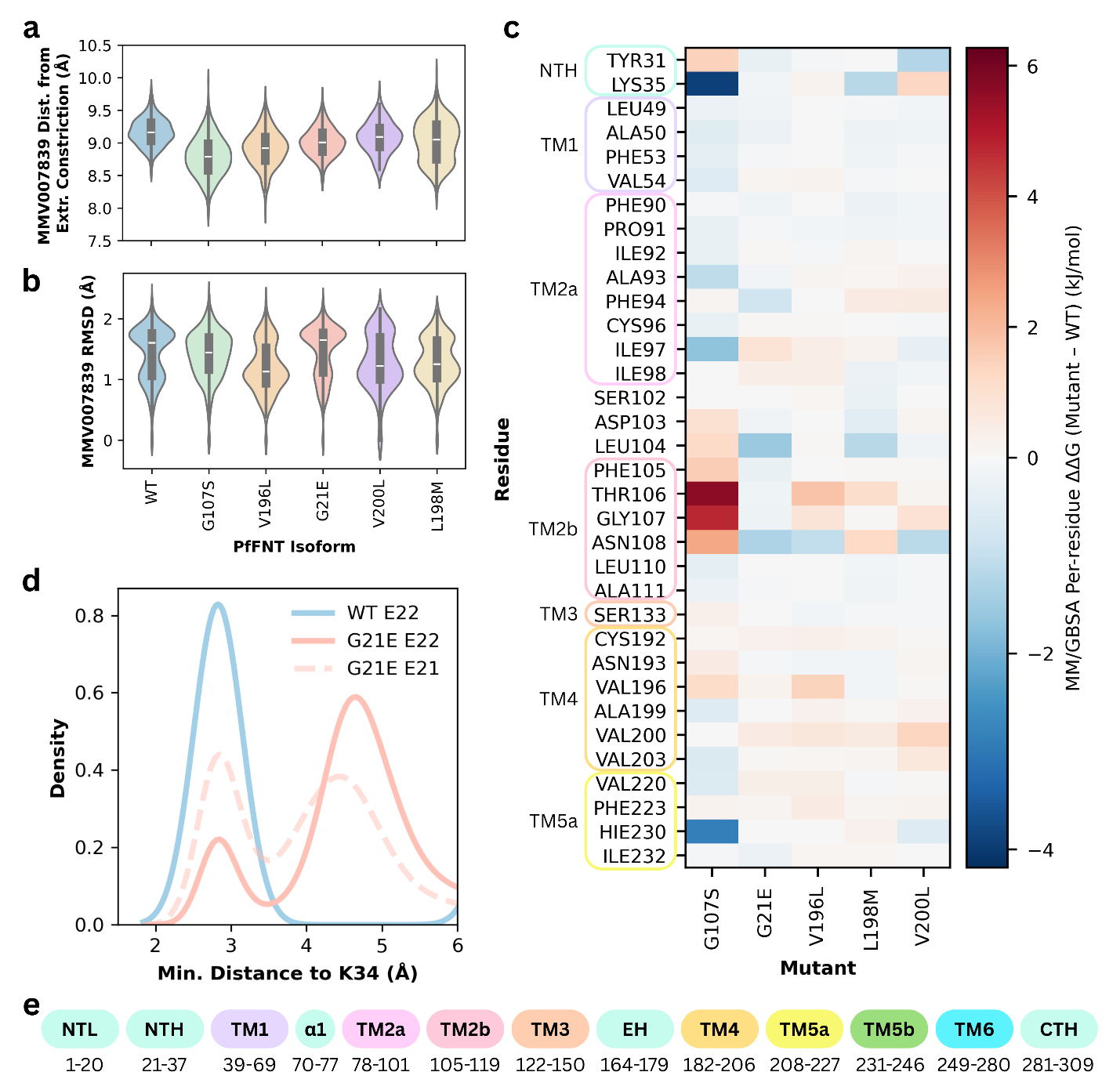

**Figure S1. Effects of known mutations in MD simulations. a)** Distance between the COM of MMV007839 and the COM of the extracellular constriction (Exr. Constr.) and **b)** RMSD of MMV007839 across WT, G107S, V196L, G21E, V200L, and L198M simulation systems. **c)** Per residue ΔΔG values from MM/GBSA calculations. The ΔΔG values were calculated by taking the difference in interaction energy between mutant and wildtype systems. **d)** Density distributions of the minimum distance between K34 and E22 in WT simulation systems (blue), K34 and E22 in G21E simulation systems (red, solid line), and K34 and E21 in G21E simulation systems. **e)** Residue ranges of major helices and loops within the PfFNT subunit.

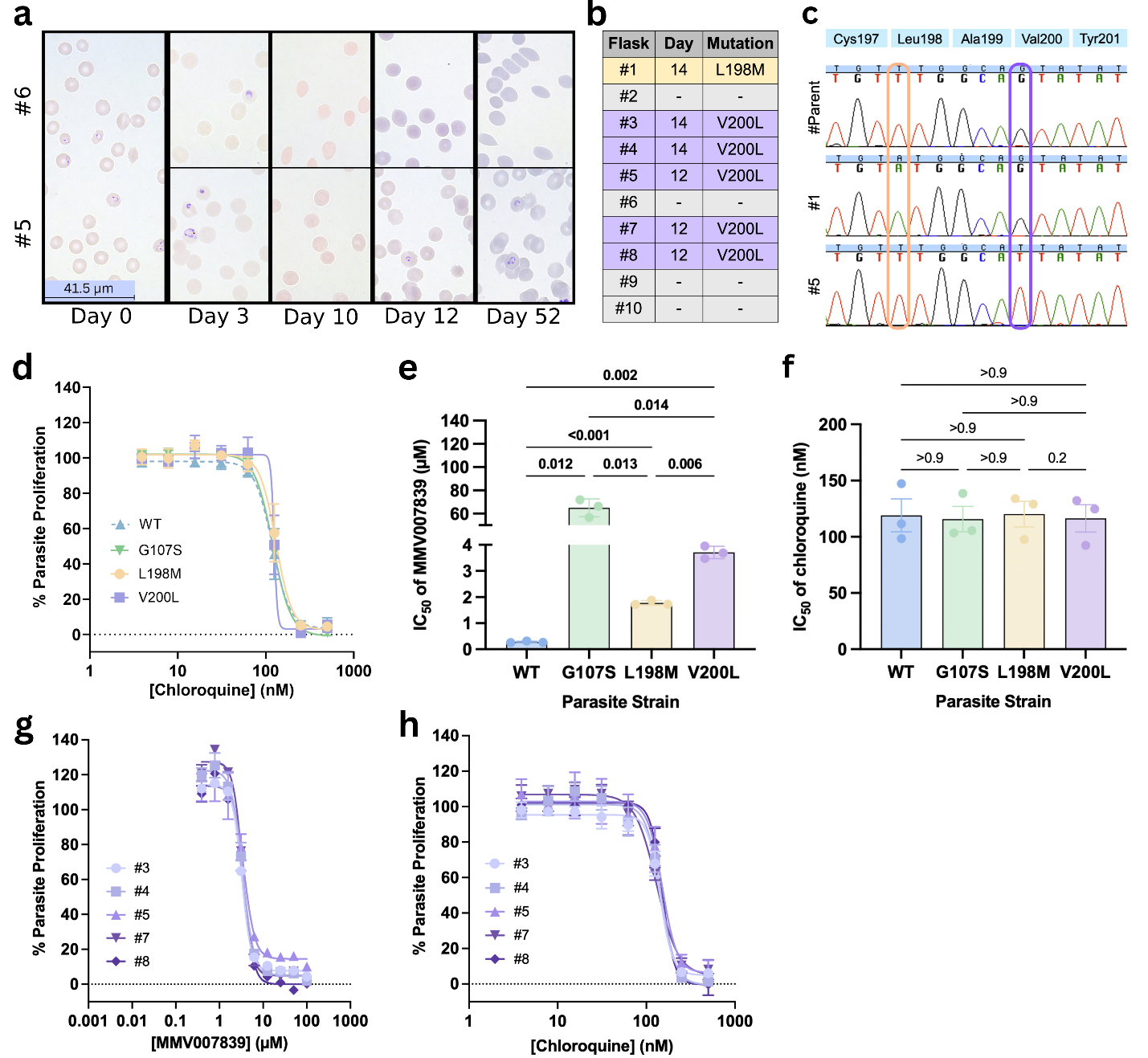

**Figure S2. The L198M and V200L mutations are associated with parasite resistance to MMV007839 but not chloroquine. a)** Images from Day 0, 3, 10, 12 and 52 of cells from a flask in which resistant parasites were generated (#5) and a flask in which no resistant parasites emerged (flask 6) during the *in vitro* evolution experiment. **b)** The first day that parasites were observed again in each of the 10 flasks, where “-” means no parasites were seen by the end of the experiment (day 52). The corresponding PfFNT mutation that was discovered in parasites from each flask is shown. **c)** Sequencing of the *pffnt* gene revealed a mutation coding for a L198M change in PfFNT in parasites from flask #1, and a mutation coding for a V200L change in PfFNT in parasites from the other five flasks in which resistant parasites emerged (data shown for flask #5). These mutations were not observed in the *pffnt* gene of the parental parasites. The single point mutation responsible for the amino acid change is highlighted in a box, and the WT amino acid is shown in a blue box above the chromatogram. **d)** The effect of chloroquine on the proliferation of Dd2^PfFNT-WT^, Dd2^PfFNT-L198M^, Dd2^PfFNT-V200L^ (flask #4) and Dd2^PfFNT-G107S^ (from Hapuarachchi et al.^3^) parasites. The data are the mean ± SEM from three independent experiments (with the exception of the highest concentration for Dd2^PfFNT-WT^, which was n = 2; each performed with internal triplicates). **e)** IC_50_ values for MMV007839 from the experiments for which data are shown in **Fig. 3a**. **f)** IC_50_ values for chloroquine from the experiments for which data are shown in **d)**. In **e)** and **f)** the symbols show the IC_50_ values obtained in individual experiments, the bars show the mean, and the error bars show the SEM. Statistical significance was calculated using a one-way ANOVA with post hoc Tukey test. Significant values are shown in bold. **g)** and **h)** The response to MMV007839 (**g**) or chloroquine (**h**) of parasites harbouring PfFNT-V200L obtained from flasks #3, #4, #5, #7 and #8. The data are from a single experiment. The error bars show the standard deviation from three technical replicates.

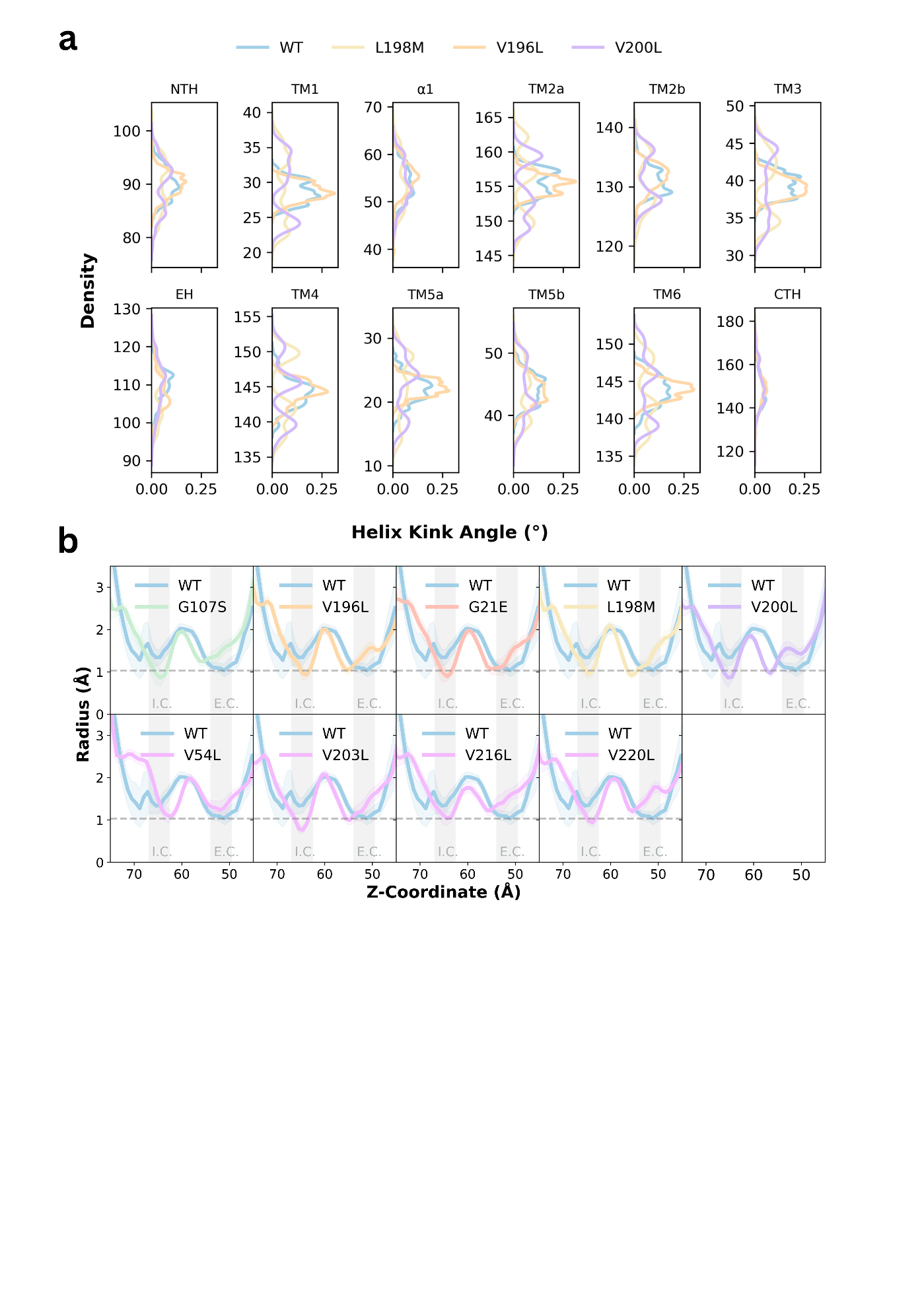

**Figure S3. Effects of mutations on helix kink angles and transport pathway radius. a)** Density plots of the kink angles (°) adopted by the NTH, TM1, α1, TM2a, TM2b, TM3, EH, TM4, TM5a, TM5b, TM6, and CTH helices across simulations from PfFNT-WT (blue), PfFNT-V196L (orange), PfFNT-L198M (yellow), and PfFNT-V200L (purple) simulations. Data were combined from each subunit from three replicates across 1 µs of simulation per replicate. **b)** Transport pathway radius measurements along the Z-coordinate from G107S, V196L, G21E, L198M, V200L, V54L, V203L, V216L, and V220L mutant systems, relative to PfFNT-WT. Data were combined from each subunit from three replicates across 1 µs of simulation per replicate. PfFNT-WT data from Wallis et al.^1^ The locations of the intracellular constriction (I.C.) and extracellular constriction (E.C.) are shown for reference.

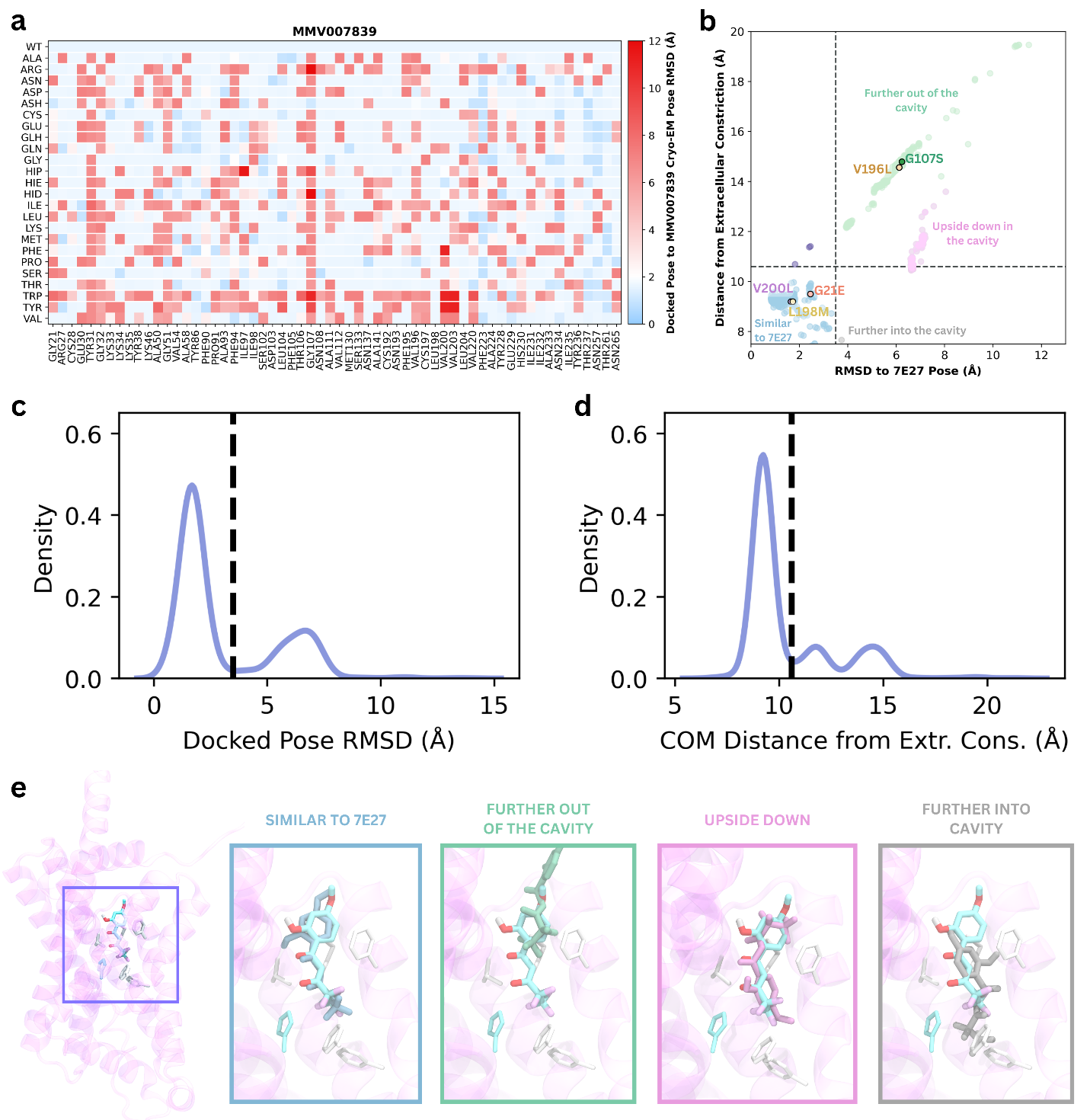

**Figure S4. Mutations identified as likely to be destabilising from docking screens.** **a)** Heatmap of the RMSD values of a representative structure from the top ranked docking pose cluster relative to the position in the 7E27 cryo-EM structure for different mutants screened in docking. **b)** Scatterplot of the RMSD values and distance from the extracellular constriction for all mutants screened in docking. The known resistance-conferring mutations are shown on the plot. The horizontal dashed line indicates the distance between the COM of MMV007839 and extracellular constriction in the 7E27 structure. The vertical dashed line at 3.5 Å RMSD is used to indicate poses that are considered destabilised relative to the pose in the 7E27 structure. Distributions of **c)** docked pose RMSD values and **d)** distances of the COM of MMV007839 in the docked pose relative to the COM of the extracellular constriction. **e)** Representative snapshots of docking poses that were either in a similar position to the pose in the 7E27 structure, further out of the cavity, upside-down within the cavity, or further into the cavity. The resolved position of MMV007839 in the 7E27 cryo-EM structure is shown in cyan in each case, and the docked pose from the mutant docking is shown in grey. Mutant poses were taken from the V200L (similar to 7E27), G107S (further out of the cavity), Y31A (upside down), and N234L (further into the cavity) docking.

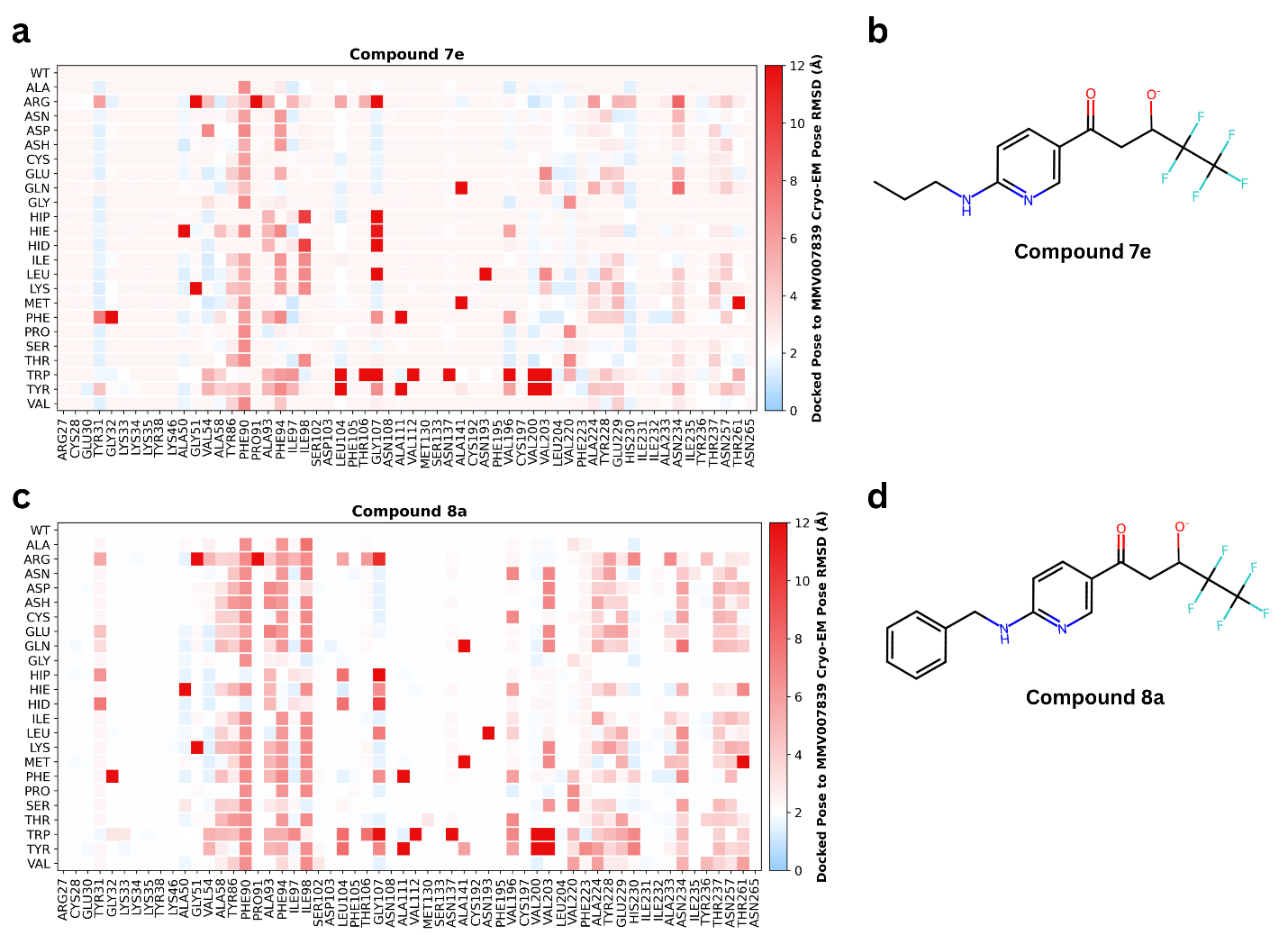

**Figure S5. Docking pipeline results for newly identified PfFNT inhibitors compounds 7e and 8a.** Heatmap of the RMSD values of a representative structure from the top ranked docking pose cluster relative to the position in the 7E27 cryo-EM structure for different mutants screened in docking for **a)** compound 7e and **c)** compound 8a. Chemical structures of **b)** compound 7e, and **d)** compound 8a.

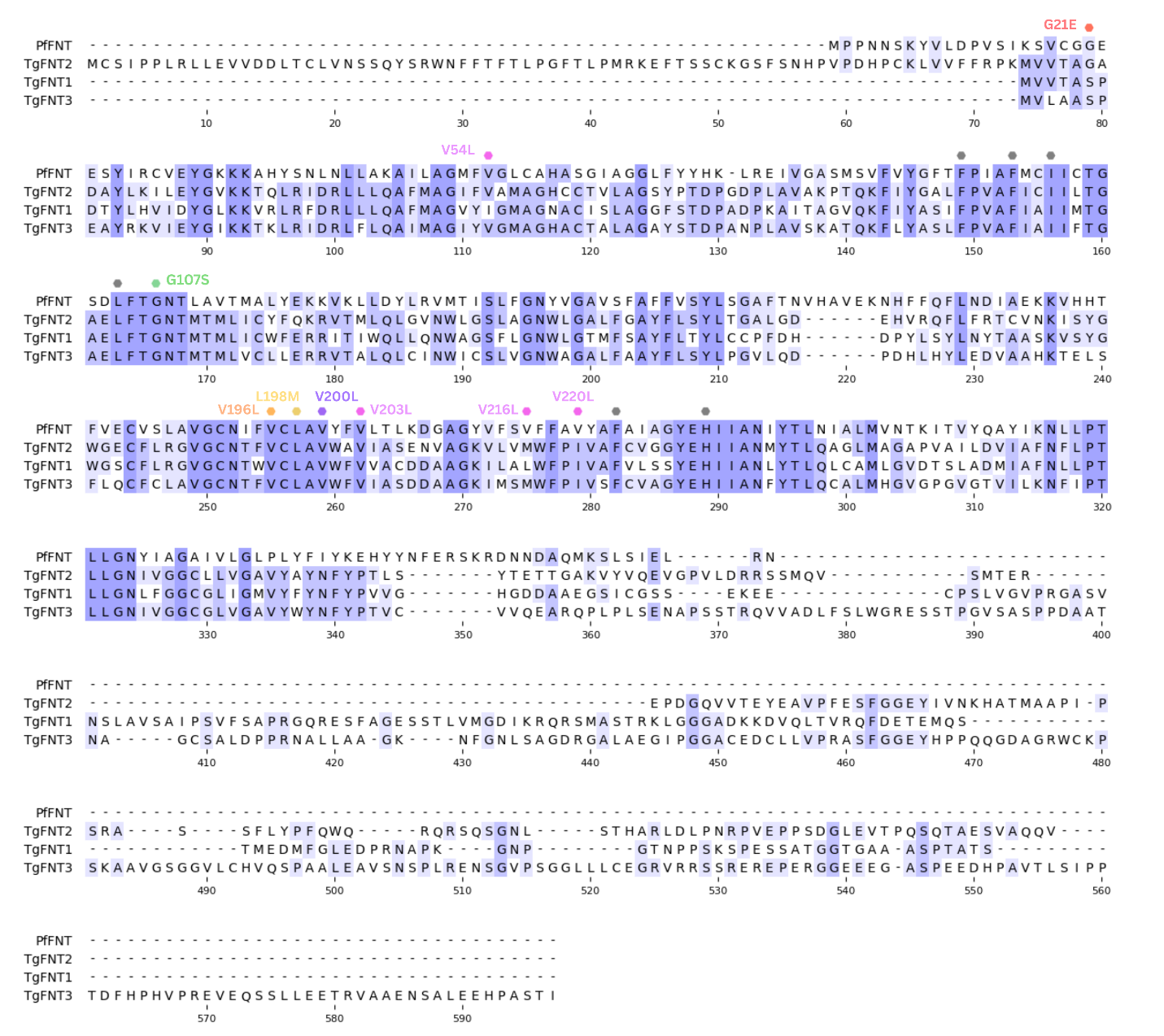

**Figure S6. Sequence alignments of PfFNT and TgFNT1, TgFNT2, and TgFNT3.** Sequence alignments for PfFNT (PF3D7_0316600 and the three *Toxoplasma gondii* FNTs, TgFNT1 (TGGT1_209800), TgFNT2 (TGGT1_292110) and TgFNT3 (TGGT1_229170). Markers indicate the locations of known resistance mutations (G107S, V196L, G21E, V200L, and L198M) and mutations along the transport pathway (V54L, V203L, V216L, and V220L). Constriction residues are marked in grey. Sequences are coloured by percent identity from lowest (white) to highest (purple). Alignment was made using Clustal Omega^4,5^ and visualised using JalView^2^.

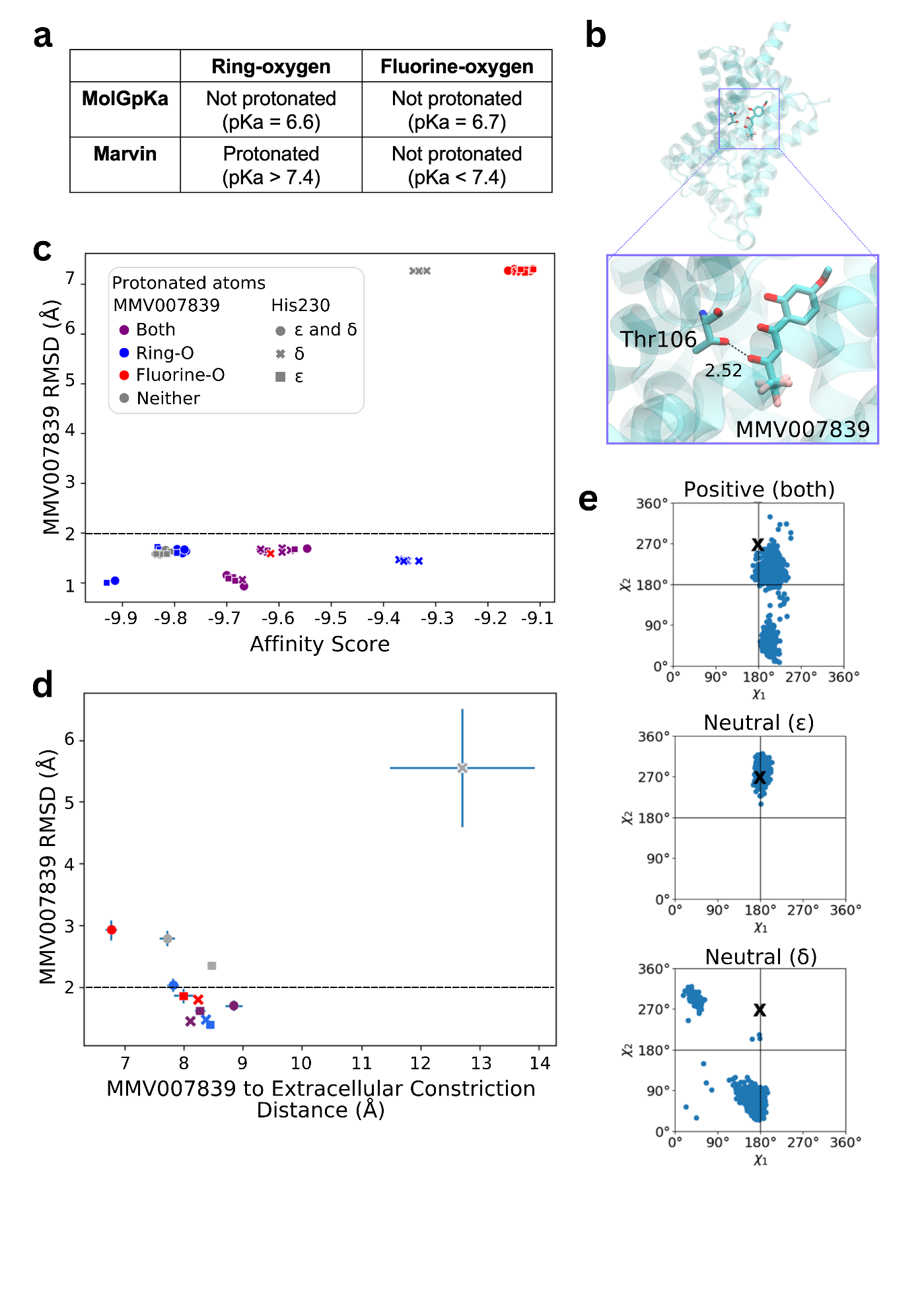

**Figure S7. Computational analysis suggests negatively charged MMV007839 binds to neutral H230. a)** pKa predictions using MolGpKa and MarvinSketch for the ring-oxygen and fluorine-oxygen of MMV007839. **b)** The cryo-EM structure of PfFNT with MMV007839 bound shows a close (2.52 Å) interaction between the fluorine-oxygen of MMV007839 and the hydroxyl oxygen of Thr106, suggesting the presence of an intermediate hydrogen atom. **c)** The RMSD change (Å) and docking affinity score (kcal/mol) of MMV007839 from docked poses for different protonation state combinations. RMSD was measured relative to the position of MMV007839 in the cryo-EM structure. Data are from five pseudo-replicates. The horizontal line shows 2 Å RMSD for reference. **d)** The position of MMV007839 from 100 ns of MD simulations at 330 K for different protonation state combinations. The RMSD of MMV007839 at the end of the simulation was calculated relative to its cryo-EM position, and was plotted against the distance of MMV007839 from the extracellular constriction. The key is shown in Panel **c**. The horizontal line shows 2 Å RMSD for reference. **e)** χ1 and χ2 sidechain dihedral angles of H230 from simulations of different His230 protonation states where MMV007839 was protonated only at the ring oxygen. Each blue dot is from every 0.33 ns of simulation. The black crosses show the starting χ1 and χ2 angles of H230 from the 7E27 cryo-EM structure.

^26^
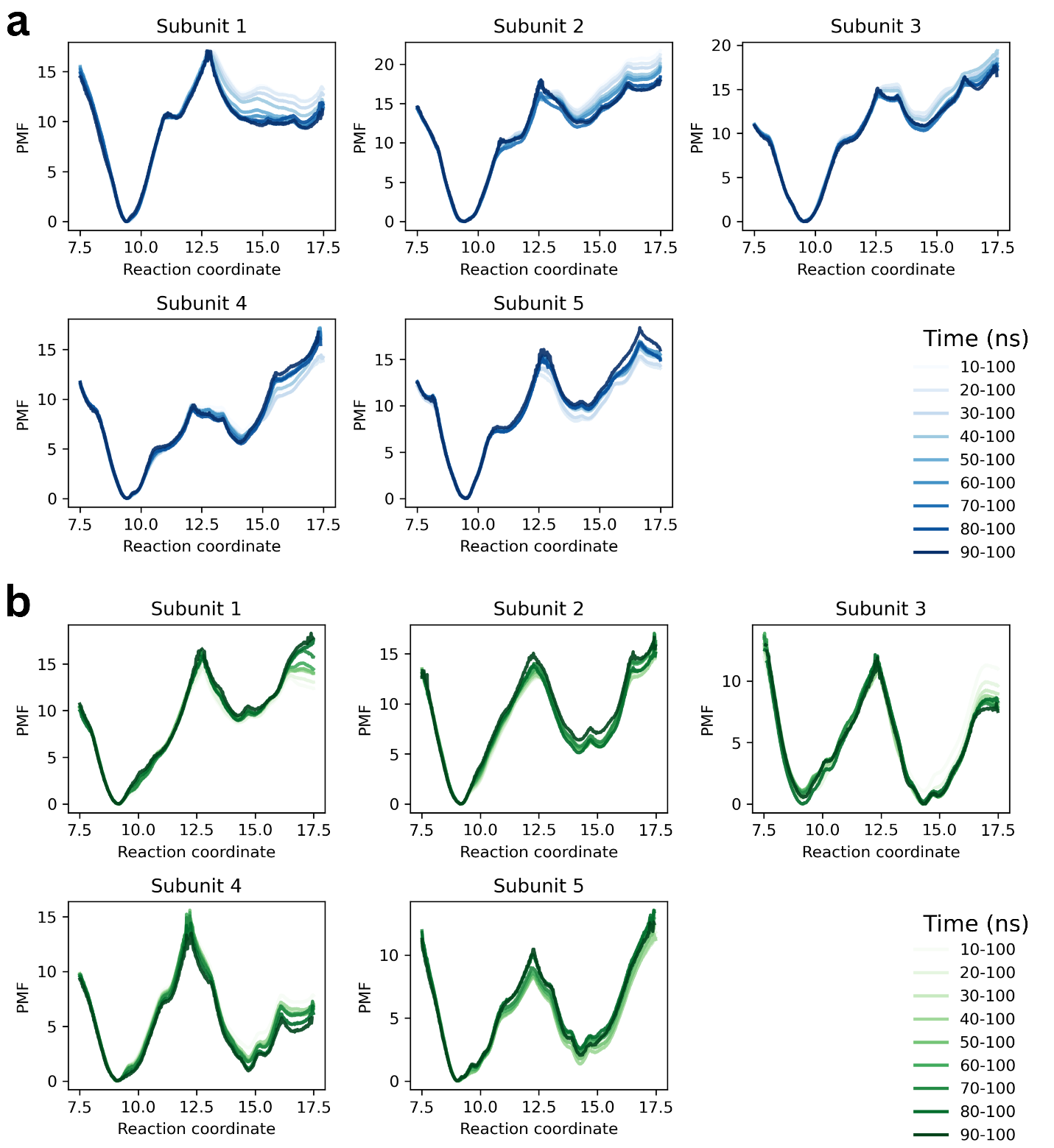

**Figure S8. PfFNT-WT and PfFNT-G107S umbrella sampling equilibration time analysis.** PMFs for each of the five subunits from **a)** PfFNT-WT and **b)** PfFNT-G107S 1D umbrella sampling simulations, constructed using different simulation lengths, are overlaid to assess the time required for equilibration. The y-axis shows the potential of mean force (PMF, kJ/mol) as MMV007839 moves through the protein. The x-axis shows the distance from the extracellular constriction site (Å) as MMV007839 moves from bulk solution into the transport cavity**.**

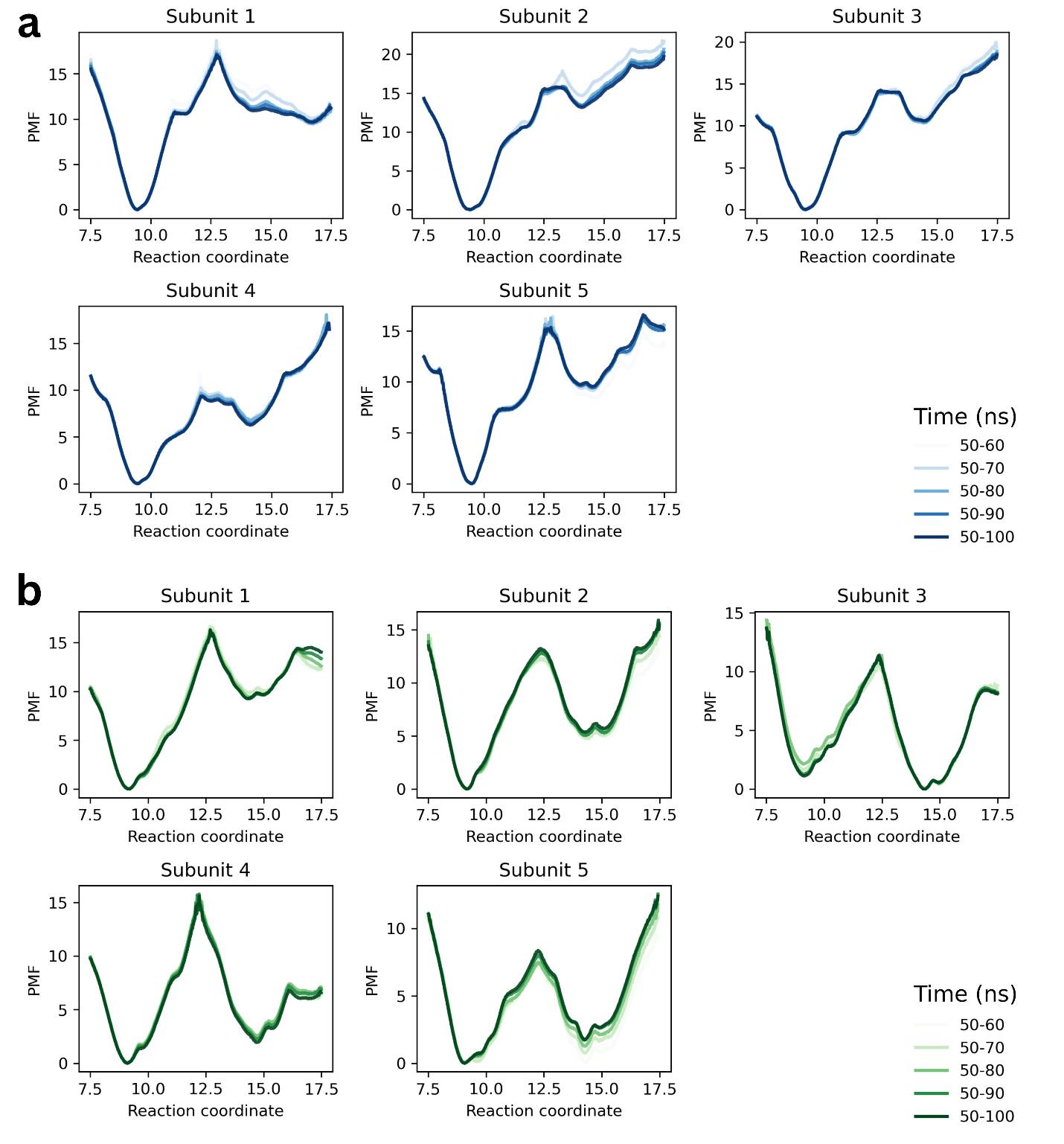

**Figure S9. PfFNT-WT and PfFNT-G107S umbrella sampling convergence analysis.** PMFs for each of the five subunits from **a)** PfFNT-WT and **b)** PfFNT-G107S 1D umbrella sampling simulations, constructed using increasing amounts of post-equilibration simulation time, are overlaid to assess convergence. The y-axis shows the potential of mean force (PMF, kJ/mol) as MMV007839 moves through the protein. The x-axis shows the distance from the extracellular constriction site (Å) as MMV007839 moves from bulk solution into the transport cavity.

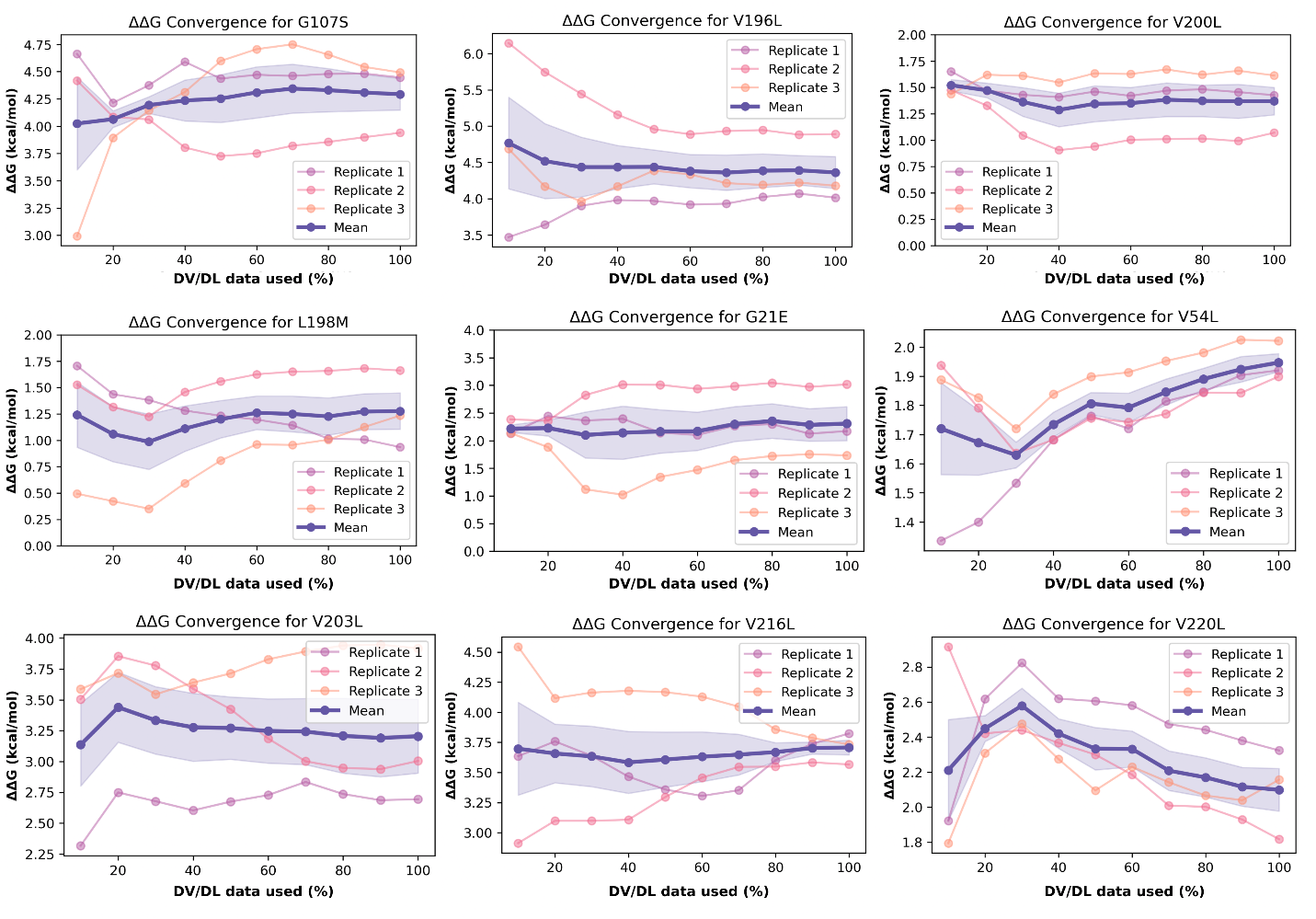

**Figure S10. Assessment of TI convergence for PfFNT mutants.** ΔΔG values were recalculated using progressively larger portions of data from each λ-window, beginning with the first 10% of DV/DL data and increasing in 10% increments up to 100%, to assess convergence of the free energy estimates. Individual replicate calculations are shown alongside the mean ± SEM.

**References**

1.	Wallis, C. *et al.* A proton transfer mechanism in the malaria parasite lactate/H+ symporter reveals a channel-like transporter without conformational changes. *bioRxiv* 2026.02.05.704099 (2026) doi:10.64898/2026.02.05.704099.

2.	Waterhouse, A. M., Procter, J. B., Martin, D. M. A., Clamp, M. & Barton, G. J. Jalview Version 2-A multiple sequence alignment editor and analysis workbench. *Bioinformatics* **25**, 1189–1191 (2009).

3.	Hapuarachchi, S. V. *et al.* The Malaria Parasite’s Lactate Transporter PfFNT Is the Target of Antiplasmodial Compounds Identified in Whole Cell Phenotypic Screens. *PLoS Pathog.* **13**, 1–24 (2017).

4.	Goujon, M. *et al.* A new bioinformatics analysis tools framework at EMBL–EBI. *Nucleic Acids Res.* **38**, W695–W699 (2010).

5.	Sievers, F. *et al.* Fast, scalable generation of high-quality protein multiple sequence alignments using Clustal Omega. *Mol. Syst. Biol.* **7**, 539 (2011).
